# Super interactive promoters provide insight into cell type-specific regulatory networks in blood lineage cell types

**DOI:** 10.1101/2021.03.15.435494

**Authors:** Taylor M. Lagler, Yuchen Yang, Yuriko Harigaya, Vijay G. Sankaran, Ming Hu, Alexander P. Reiner, Laura M. Raffield, Jia Wen, Yun Li

**Affiliations:** Department of Biostatistics, University of North Carolina at Chapel Hill, Chapel Hill, NC 27599, USA; Department of Pathology and Laboratory Medicine, University of North Carolina at Chapel Hill, Chapel Hill, NC, 27599, USA; McAllister Heart Institute, University of North Carolina at Chapel Hill, Chapel Hill, NC, 27599, USA; Curriculum in Bioinformatics and Computational Biology, Department of Genetics, University of North Carolina at Chapel Hill, Chapel Hill, NC 27599, USA; Division of Hematology/Oncology, Boston Children’s Hospital and Department of Pediatric Oncology, Dana-Farber Cancer Institute, Harvard Medical School, Boston, MA 02115, USA; Broad Institute of MIT and Harvard, Cambridge, MA 02142, USA; Harvard Stem Cell Institute, Cambridge, MA 02138, USA; Department of Quantitative Health Sciences, Lerner Research Institute, Cleveland Clinic Foundation, Cleveland, Ohio 44195, USA; Division of Public Health Sciences, Fred Hutchinson Cancer Research Center, Seattle, WA 98109, USA; Department of Epidemiology, University of Washington, Seattle, WA 98195, USA; Department of Genetics, University of North Carolina at Chapel Hill, Chapel Hill, NC 27514, USA; Department of Computer Science, University of North Carolina at Chapel Hill, Chapel Hill, NC 27599, USA

## Abstract

Existing studies of chromatin conformation have primarily focused on potential enhancers interacting with gene promoters. By contrast, the interactivity of promoters *per se*, while equally critical to understanding transcriptional control, has been largely unexplored, particularly in a cell type-specific manner for blood lineage cell types. In this study, we leverage promoter capture Hi-C data across a compendium of blood lineage cell types to identify and characterize cell type-specific super-interactive promoters (SIPs). Notably, promoter-interacting regions (PIRs) of SIPs are more likely to overlap with cell type-specific ATAC-seq peaks and GWAS variants for relevant blood cell traits than PIRs of non-SIPs. Further, SIP genes tend to express at a higher level in the corresponding cell type, and show enriched heritability of relevant blood cell trait(s). Importantly, this analysis shows the potential of using promoter-centric analyses of chromatin spatial organization data to identify biologically important genes and their regulatory regions.

## Background

Genome-wide chromosome conformation capture techniques such as Hi-C [1] have been widely used to study chromatin three dimensional (3D) organization. However, due to the complexity and sparsity of Hi-C data, it is difficult to identify statistically significant long-range chromatin interactions between distant genomic sequences at fine resolutions (e.g, at restriction fragment level, or < 10Kb equal size bin level) even with tens of billions of pairwise reads produced [2, 3]. Furthermore, ultra-deep sequencing is costly and likely to generate redundant reads, leading to Hi-C library saturation [4]. In addition, chromatin spatial organization studies have largely focused on regulatory regions, but characterization of the 3D genome at promoters is also important for understanding gene expression regulation. To bridge this gap, capture Hi-C and subsequent variations were developed as an extension of the Hi-C technique by combining target enrichment and sequencing [5–8]. One such capture technique, promoter capture Hi-C (pcHi-C), was developed to focus on promoter regions. These regions have been largely taken for granted and automatically removed from detailed study in many chromatin conformation-based studies [9–11]. pcHi-C is specifically enriched for promoter sequences and enables the genome-wide detection of distal promoter-interacting regions (PIRs), for all promoters with a priori designed probes/baits in a single experiment [12].

Promoter interactomes (the set of all interactions involving promoters within a cell) are tissue-and lineage-specific and have been used to link promoters to GWAS risk loci [9, 12–14]. Consequently, there has been growing interest in studying cell type-specific differences in PIRs. As one example, pcHi-C analysis of 17 human hematopoietic cells demonstrated that PIRs are highly cell type-specific and reflective of the expected lineage relationships (such as mapping of promoter interactions for T-cell receptor component encoding genes to lymphoid cell types only, not to myeloid lineage cell types). Importantly, this analysis demonstrated the ability of pcHi-C to link non-coding regulatory variants to their target genes [13]. Thus, pcHi-C analysis can be leveraged to provide insight into gene expression control and the function of non-coding disease-associated sequence variants [12].

A recent study on human corticogenesis has identified a subset of promoters exhibiting unusually high degrees of chromatin interactivity (where chromatin interactivity is defined by cumulative CHiCAGO scores of interactions with neighboring regions), which were termed super-interactive promoters (SIPs) [8]. Song et al. found that these brain cortex SIPs were enriched for corresponding lineage-specific genes, suggesting that the interactions between SIPs and their regulatory networks may play a role in modulating cell type-specific transcription. In addition, Song et al. also found SIPs in hematopoietic lineages using pcHi-C data, but did not perform further annotation or characterization of these hematopoietic SIPs.

Due to the relative ease of measuring blood cells, rich genomics data is available for hematopoietic cells. Further, different hematopoietic cell types play different roles in blood cell generation and function and correspond to different phenotypic traits (for example inflammation, autoimmunity, and infection phenotypes for white blood cell types, thrombosis and hemostasis related phenotypes for platelet producing megakaryocytes), emphasizing the importance of studying them in cell type-specific manner [15]. Blood cells are highly relevant tissues for many complex phenotypes, including infectious disease susceptibility (including COVID-19), disease related biomarkers such as telomere length or circulating inflammatory cytokines, thrombosis (including venous thromboembolism and stroke), asthma and other respiratory diseases, and autoimmune conditions [16]. Understanding of interactions of gene promoters and their regulatory regions in specific blood cell types, as opposed to simple analysis of “whole blood”, can lead to improved annotation of genome-wide association study (GWAS) identified loci and their target genes, and thus of the genetic mechanisms underlying complex disease risk. Hematopoietic SIPs are thus of broad interest for understanding gene regulation and its connection to disease risk in human populations.

Here, we focus on characterizing promoter-centric chromatin spatial interaction profiles, across a compendium of cell types in the hematopoietic lineage. In this study, we identify and characterize SIPs in human blood cells using pcHi-C data from the Javierre et al. study [13]. We find that SIPs tend to be cell type-specific or shared across all cell types. Through examining the differences between SIPs and non-SIPs in terms of their interaction profiles as well as their genes, we find that SIPs share common properties across cell types. Importantly, we demonstrate how studying SIP networks may provide insight into the complex regulation of promoters as well as potential functional interactions.

## Results

### Cell Type-Specifically Expressed Genes Exhibit Higher Levels of Chromatin Interactivity in the Corresponding Cell Type than Shared Genes

We first explored the relationship between chromatin interactivity and gene expression in a cell type-specific manner. We examined this relationship using pcHi-C data from Javierre et al. [13] and gene expression data from BLUEPRINT [17], in each of the five hematopoietic cell types: erythrocyte (Ery), macrophage/monocyte (MacMon), megakaryocyte (MK), naive CD4 T-cell (nCD4), and neutrophil (Neu) (Methods). We classified genes as “specific” (expressed in a cell type-specific manner) or “shared” across the five cell types (Methods). The promoters for cell type-specific genes have significantly more interactions than the shared genes across all five cell types (*p*-value < 0.05) (Figure S1a-e). Similar results were observed by Song et al. in neuron cells [8].

### Inequality in the Promoter Interactome: Few Super-Interactive Promoters

For each cell type, we ranked the promoter-containing anchor bins (baits) according to their cumulative interaction scores (Methods) (Figure 1). We find that a small number of promoter baits (∼7.5%) have extremely high cumulative interaction scores, as defined based on the curve inflection point in each cell type, and annotated them as super-interactive promoters (SIPs). In total, we annotate 1,157, 808, 1,287, 993, and 861 SIPs in erythrocytes, macrophages/monocytes, megakaryocytes, naive nCD4 T-cells, and neutrophils, respectively (Table S1). These SIPs can be cell type-specific or shared across cell types. There are 170 SIPs shared across all five cell types, as well as 189, 107, 302, 283, and 274 cell type-specific SIPs in erythrocytes, macrophages/monocytes, megakaryocytes, naive nCD4 T-cells, and neutrophils, respectively. Figure S2 details how the SIPs are shared across the different cell types (Additional File 1). Similar to GTEx analyses of eQTLs’ tissue specificity [18, 19], the most common configurations pertain to cell type-specific SIPs and shared SIPs (across all five cell types). In addition, principal component analysis (PCA) on the cumulative interaction scores reflects expected correlations between cell type-specific SIPs in each cell type, as well as between any SIP and those SIPs shared by all five cell types (Figure S3).

**Figure 1.**
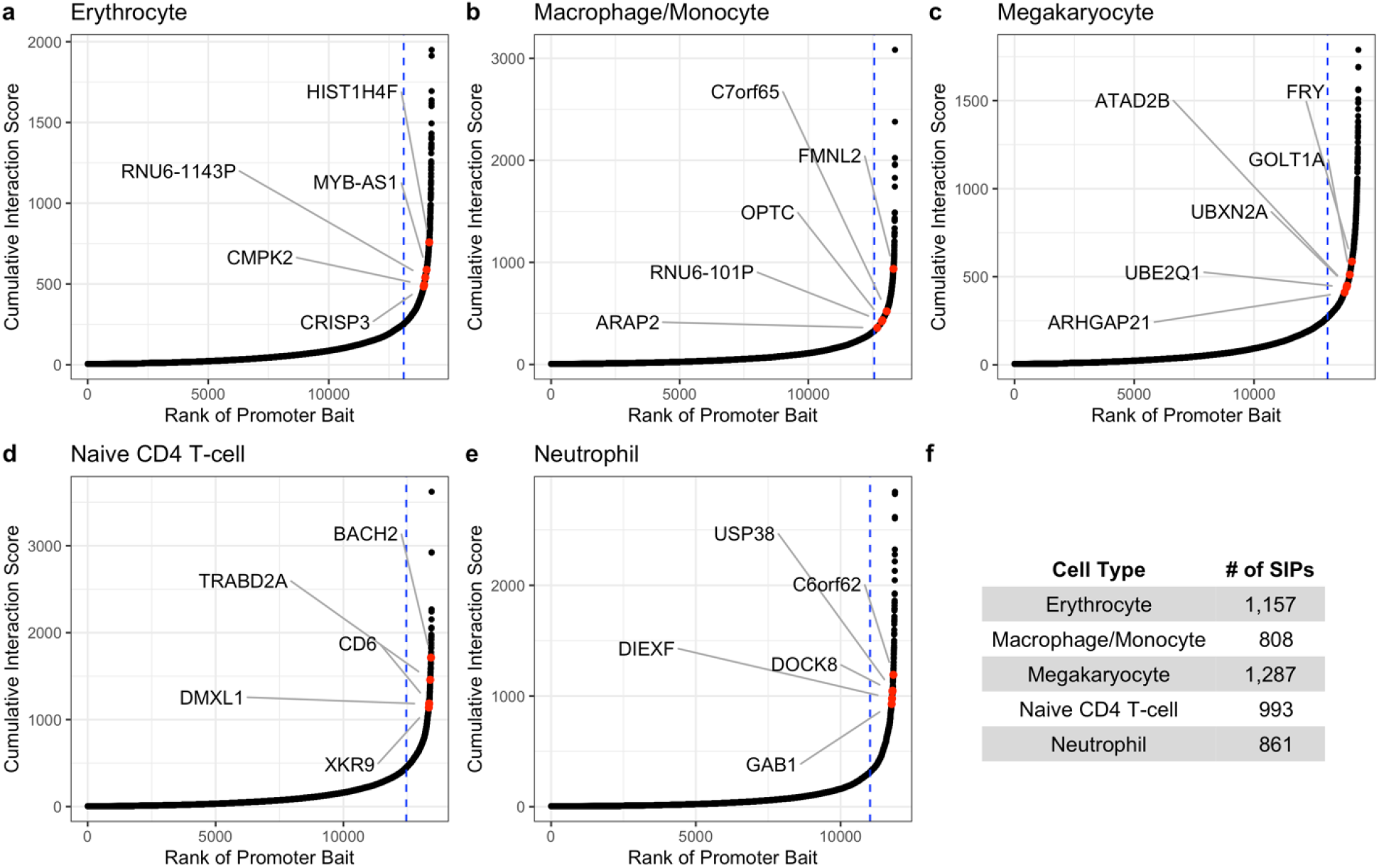
There are few super-interactive promoters (SIPs) in the chromatin interactome. **(a)-(e)** Hockey plots for each cell type show the ranked cumulative interaction scores for pcHi-C promoter-containing anchor bins (baits). Promoters to the right of the blue vertical line are classified as super interactive promoters (SIPs), as they exhibit unusually high levels of chromatin interactivity. Red dots highlight the highest ranked cell type-specific SIP genes^1^ with a PIR overlapping a relevant GWAS identified SNP. **(f)** Total number of SIPs annotated per cell type. 1 The megakaryocyte genes *ATAD2B* and *UBXN2A* correspond to the same SIP bait.

Moreover, many cell type-specific SIPs correspond to known lineage-specific genes and have PIRs overlapping relevant GWAS variants (examples annotated by red dots in Figure 1a-e) (Methods, Additional File 2). For example, the neutrophil SIP gene *DOCK8* is an immunodeficiency gene that is expressed in resting human neutrophils [20], and the macrophage SIP gene *FMNL2* is most highly expressed in macrophages and is cell type relevant [21–23]. The naive CD4 T-cell SIP gene *CD6* is a strong positive control, as this gene is essentially only expressed in CD4 T-cells [24]; *BACH2* plays a vital role in maintaining naive CD4 T-cells and regulating immune homeostasis [25]. All of these SIP genes have at least one PIR overlapping a GWAS identified SNP.

The unusually high cumulative interaction scores at SIPs are driven by a large number of interactions, rather than a few interactions with large scores (Figure 2a-b). SIP baits have a significantly greater number of other end interactions (i.e., PIRs) compared to non-SIP baits in each cell type (Wilcoxon *p*-value < 2.2e-16). The median number of significant interactions is 38-61 for SIPs and only 4-7 for non-SIPs. SIPs interact with ∼9 times more PIRs than non-SIPs on average. However, the median CHiCAGO score [26] of significant interactions per bait, although statistically different, is comparable between SIPs and non-SIPs (the median is ∼8.4 for SIPs and ∼6.4 for non-SIPs).

**Figure 2.**
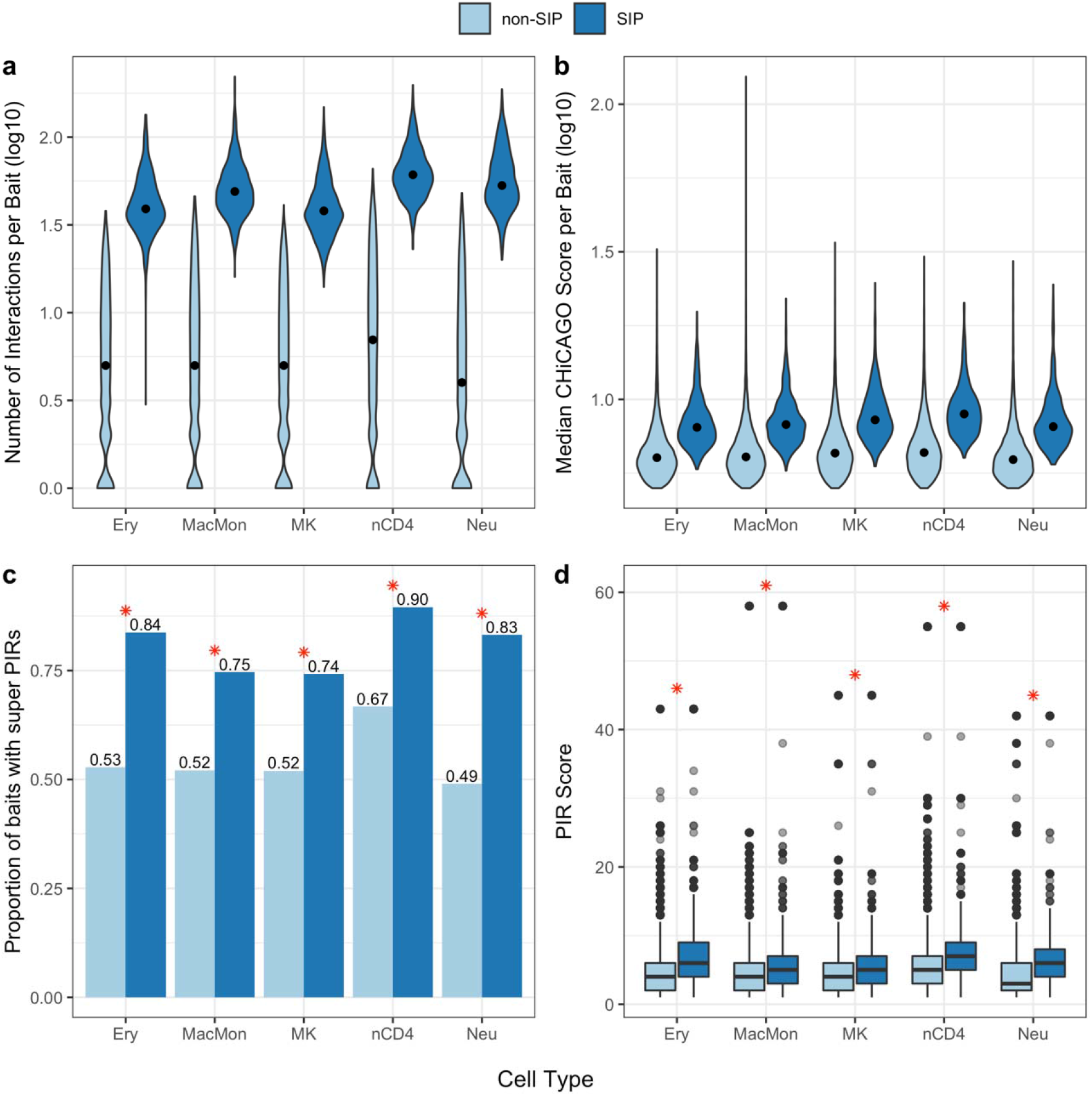
SIPs are driven by a large number of interactions. **(a)** Distribution of the number of significant interactions (log10 scale) between promoter bait and promoter interacting regions (PIRs) for SIPs and non-SIPs in each cell type. **(b)** Distribution of the median CHiCAGO score (log10 scale) of significant interactions per promoter bait for SIPs and non-SIPs in each cell type. The width of each violin corresponds to the frequency of interaction count (a) or median CHiCAGO score (b). The median of each distribution is marked by a black dot. **(c)** The proportion of SIP and non-SIP baits with super PIRs in each cell type. SIPs interact with a larger proportion of super PIRs than non-SIPs in each cell type (the red asterisk (*) denotes Chi-square *p*-value < 3.2e-35). **(d)** Distribution of PIR scores for SIPs and non-SIPs in each cell type. SIPs have significantly higher PIR scores than non-SIPs in each cell type (the red asterisk (*) denotes Wilcoxon *p*-value < 1.7e-50). Details for panels (c) and (d) can be found in Table S1. (Ery = erythrocytes; MacMon = macrophages/monocytes; MK = megakaryocytes; nCD4 = naive CD4 T-cells; Neu = neutrophils)

### SIPs and Super Promoter-Interacting Regulatory Regions

In each cell type, ∼59% of PIRs interact with a single promoter fragment while only ∼10% of PIRs interact with 4 or more promoter fragments. Therefore, we define a super promoter-interacting region (super PIR) as a PIR interacting with at least 4 promoter fragments. As expected, SIPs interact with a larger proportion of super PIRs than non-SIPs in each cell type (Chi-square *p*-value < 3.2e-35) (Figure 2c, Table S2). Approximately 74-90% of SIPs interact with a super PIR, whereas only 49-67% of non-SIPs interact with a super PIR. We assign each promoter region (bait) a PIR score, defined by its PIR with the maximum number of interactions. SIPs have significantly higher PIR scores than non-SIPs in each cell type (Wilcoxon *p*-value < 1.7e-50) (Figure 2d, Table S3). The median PIR score is ∼6 for SIPs and ∼4 for non-SIPs. The basic characteristics of SIPs (e.g., number of PIRs and proportion with super PIRs) are consistent across all five hematopoietic cell types.

### SIP PIRs Overlap with ATAC-seq Peaks and Relevant GWAS Variants

We can further characterize SIPs through their PIRs by looking at the proximity of PIRs to open chromatin regions and known GWAS variants. In each cell type, over 96% of SIPs have a PIR overlapping an ATAC-seq peak of the corresponding cell type [27], compared to 63-83% of non-SIPs. In each cell type, the proportion of SIPs with a PIR that overlaps a cell type-specific ATAC-seq peak is significantly greater than the proportion of non-SIPs with a PIR that overlaps an ATAC-seq peak (Chi-square *p*-value < 2.9e-45) (Figure 3a). We then compared the number of PIRs overlapping cell type-specific ATAC-seq peaks, for SIPs and non-SIPs (Figure 3b). In each cell type, significantly more SIP PIRs overlap with cell type-specific ATAC-seq peaks compared to non-SIP PIRs (*t*-test *p*-value < 1.2e-162). The median number of ATAC-seq overlaps per bait is 8-22 for SIPs and only 1-3 for non-SIPs. Details on the number of overlaps as well as specific *p-*values are reported in Table S3. Note that neutrophils are excluded from this analysis due to data availability (Supplemental Note 1).

**Figure 3.**
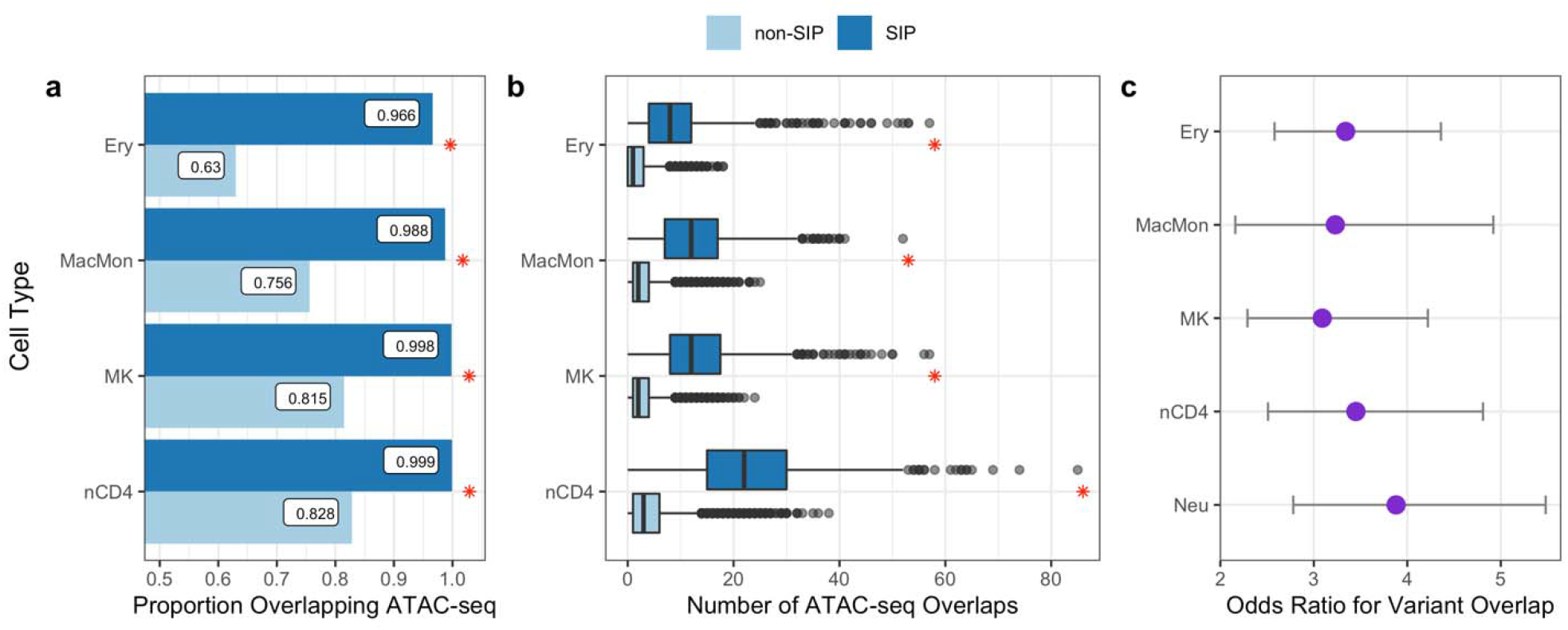
SIP PIRs overlap with ATAC-seq peaks and relevant GWAS variants. **(a)** In each cell type, the proportion of SIPs with a PIR that overlaps a cell type-specific ATAC-seq peak is significantly greater than the proportion of non-SIPs with a PIR that overlaps an ATAC-seq peak (the red asterisk (*) denotes Chi-square *p*-value < 2.2e-16). **(b)** The distribution of the number of PIRs (other ends) overlapping with ATAC-seq peaks for each pcHi-C bait (y-axis) for each cell type (x-axis). In each cell type, significantly more SIP interactions overlap with ATAC-seq peaks compared to non-SIP interactions (the red asterisk (*) denotes two-sided t-test *p*-value < 1.2e-162). **(c)** In each cell type, SIPs have 3-4 times the odds of having at least one PIR overlap with a relevant GWAS variant, compared to non-SIPs. Odds ratio estimates (purple dots) and corresponding 95% confidence intervals are shown. (Ery = erythrocytes; MacMon = macrophages/monocytes; MK = megakaryocytes; nCD4 = naive CD4 T-cells; Neu = neutrophils)

Next, we examine the overlap between GWAS variants and PIRs. In each cell type, SIPs have significantly greater odds (3-4 times the odds) of having at least one PIR overlap with a relevant blood cell trait associated variant, compared to non-SIPs (Methods) (Figure 3c). We found similar results when considering only cell type-specific SIPs (Figure S4) and observed that the basic characteristics of SIPs are consistent across all five cell types. Details SIP PIRs and their overlaps with relevant variants can be found in Additional File 2.

### SIP Subnetworks

By incorporating GWAS and open chromatin data with the pcHi-C data, we can determine SIP subnetworks that may provide insight into potential functional interactions. These SIP subnetworks are defined as having at least two PIRs that each overlap with a relevant statistically independent SNP and a cell type-specific ATAC-seq peak (Methods). We identify 2-15 SIP subnetworks in each cell type/phenotype combination (Methods, Additional File 2). Details of the interactions and SNPs involved in these SIP subnetworks can be found in Additional File 3.

We highlight two examples of SIP subnetworks in Figure 4. Figure 4a depicts the megakaryocyte SIP with bait located at the *EPHB3* gene interacting with three distinct regions that overlap with a total of 8 independent SNPs related to platelet count. These PIRs also overlap with megakaryocyte ATAC-seq peaks, and are near the key platelet related gene *THPO* or thrombopoietin, variants in which can lead to thrombocythemia (OMIM 600044 [28]). Thrombopoietin is essential for megakaryocyte proliferation and maturation, as well as for production of platelets. *EPHB3* encodes ephrin receptor B3, and plays roles in development, cell migration, and adhesion; variants in family member *EPHB2*, which also binds ephrin-B family ligands, are associated with a Mendelian bleeding disorder characterized by deficiencies in agonist-induced platelet aggregation and granule secretion (OMIM 600997 [28]). This SIP network suggests that *THPO* locus variants may also play a role in regulation of *EPHB3*. Figure 4b depicts the naive CD4 T-cell SIP with bait located at the *ETS1* gene interacting with three distinct PIRs that each overlap with an independent GWAS SNP related to lymphocyte count as well as a naive CD4 T-cell ATAC-seq peak. *ETS1* is a transcription factor highly expressed in CD4 T-cells known to regulate differentiation, survival and proliferation of lymphoid cells [17]; the *ETS1* locus is an important genetic regulator of risk for the autoimmune disorder systemic lupus erythematosus [29]. These SIP networks show the complex regulation of promoters for important hematopoietic cell type genes, with multiple distinct genetic variants and regions of open chromatin acting together to regulate genes.

**Figure 4.**
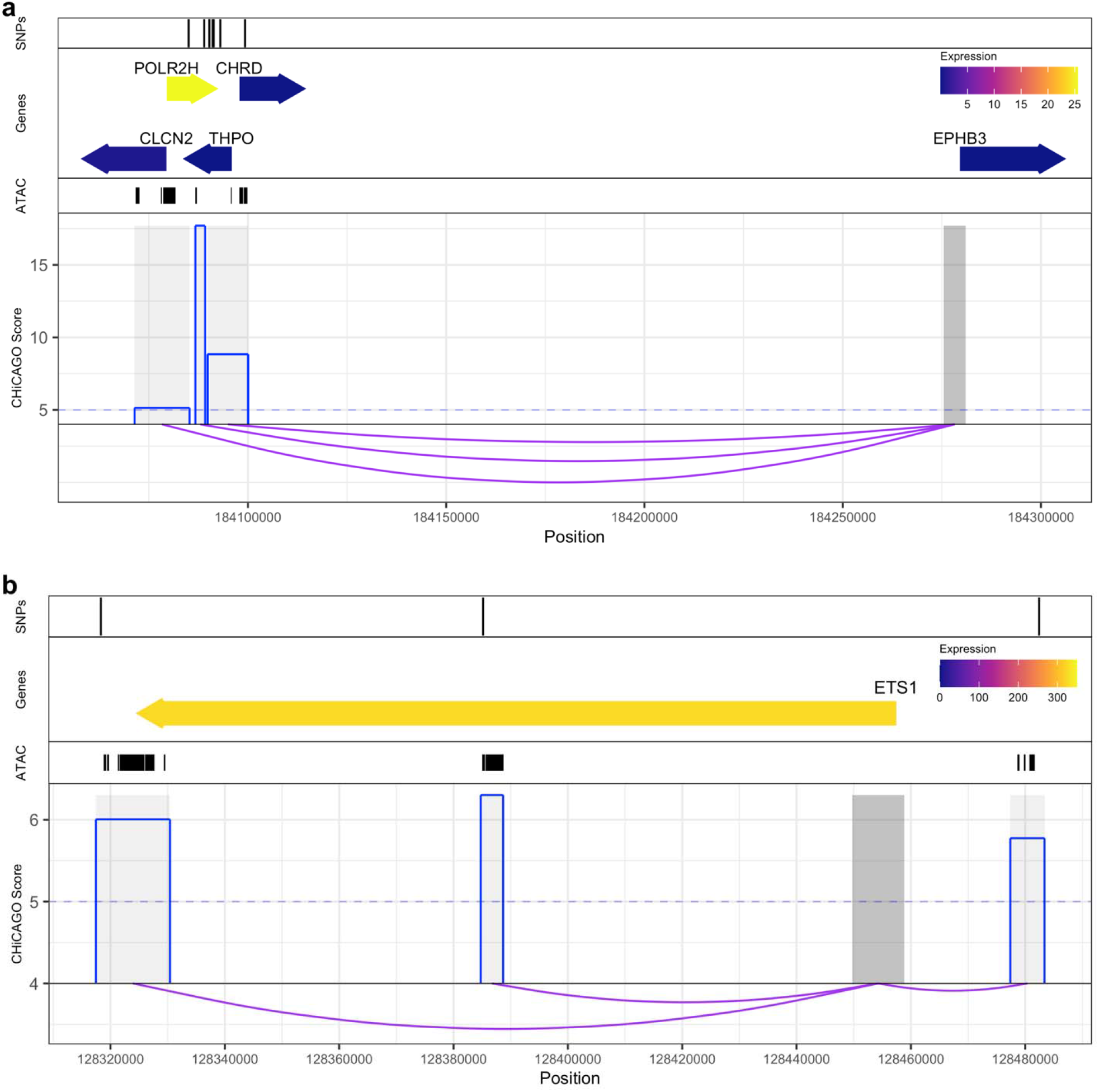
SIP subnetworks show the complex regulation of promoters for important hematopoietic cell type genes. First panel: position of SNPs. Second panel: position of SIP bait or PIR target gene(s), where the color corresponds to their exponentiated BLUEPRINT gene expression (equivalent to RKPM) in the respective cell type. Third panel: cell type-specific ATAC-seq peaks. Fourth panel: CHiCAGO scores (blue) of the interactions (depicted by purple arcs) between the SIP bait (dark grey) and the SIP PIRs (light grey). **(a)** Example of a megakaryocyte SIP subnetwork for platelet count related variants. **(b)** Example of a naive CD4 T-cell SIP subnetwork for lymphocyte count related variants.

### SIPs Align with Gene Expression Levels in a Cell Type-Specific Manner

SIPs can also be characterized by their genes, and each SIP bait may correspond to more than one gene. There are 1,514, 1,093, 1,752, 1,393, and 1,201, SIP genes in erythrocyte, macrophage/monocyte, megakaryocyte, naive CD4 T-cell, and neutrophil SIPs, respectively (Additional File 1). Within each cell type, we ranked the genes according to their expression levels and calculated the fold enrichment of the genes classified as SIPs for higher gene expression (detailed in Methods). All five cell types have well-expected trends in the relationship between SIP enrichment and gene expression (Figure 5). For example, in erythrocytes there is 1.9 fold enrichment for a gene having a SIP in the highest quintile of gene expression (1st ranked) over the lowest (5th ranked) gene expression quintile (Chi-square *p*-value = 8.7e-14).

**Figure 5.**
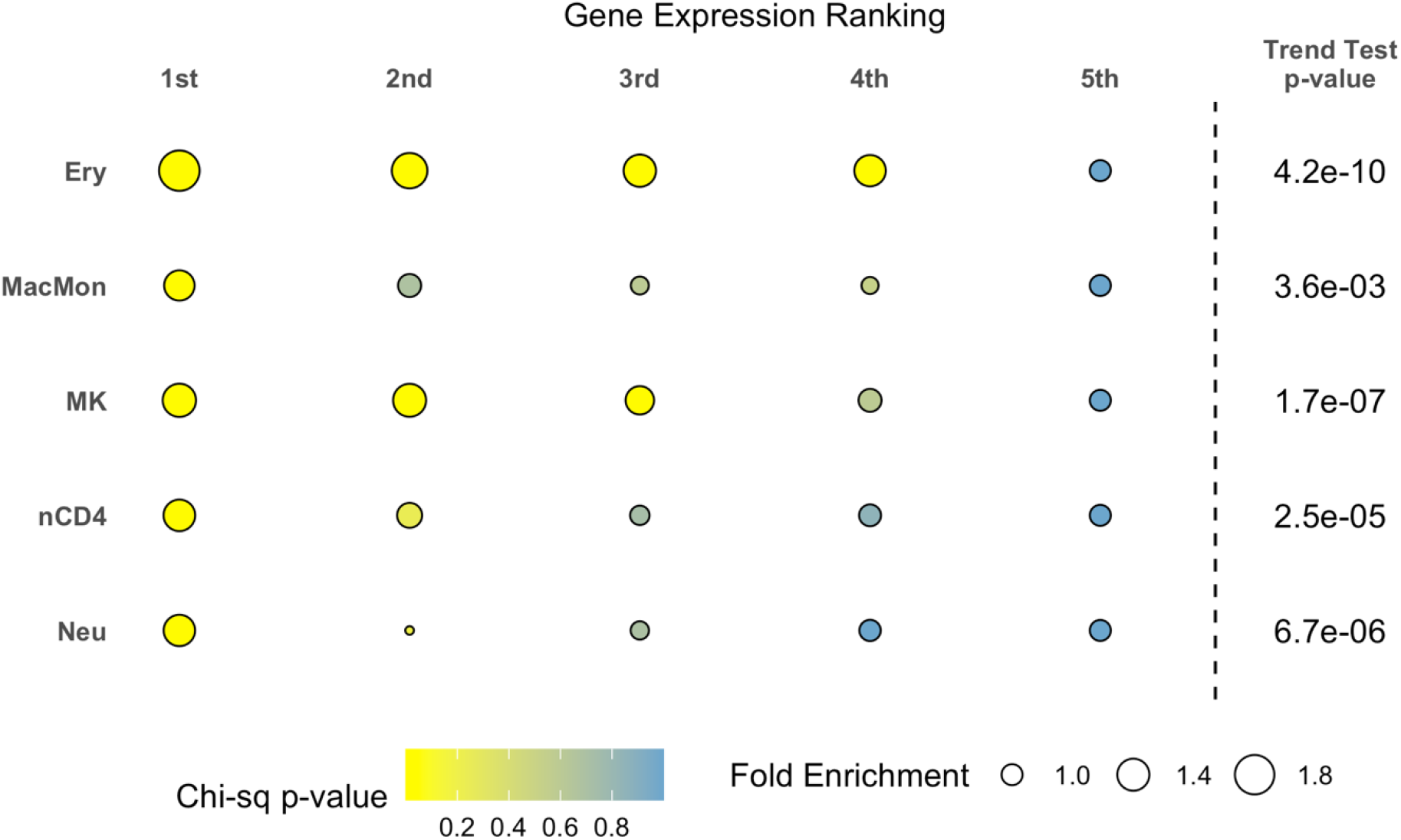
SIP genes tend to express at a higher level in the corresponding cell type. In each cell type, SIP genes with the highest (1st) ranked expression show greater fold enrichment over the lowest (5th) ranked gene expression. The size of the circle denotes the fold-change and the color denotes the Chi-square significance of enrichment. The Cochran-Armitage trend test *p*-value (one-sided) is also reported. (Ery = erythrocytes; MacMon = macrophages/monocytes; MK = megakaryocytes; nCD4 = naive CD4 T-cells; Neu = neutrophils)

### Cell Type-Specific SIP Genes

We defined cell type-specific SIP genes as genes corresponding to cell type-specific SIP baits that are not captured by any other cell type-specific or with any shared SIP baits (some genes may be captured by multiple pcHi-C baits). In total, we annotate 251, 125, 385, 386, and 384 cell type-specific genes in erythrocytes, macrophages/monocytes, megakaryocytes, naive CD4 T-cells, and neutrophils, respectively (Table S1). We also annotate 234 “shared” SIP genes (corresponding to SIPs shared across all five cell types). Note that a SIP may be a promoter for multiple genes, and thus the number of cell type-specific SIP genes may be greater than the number of cell type-specific SIPs. Full details of the SIP genes in each cell type can be found in Additional File 1.

We notice some trends in the gene expression of the 234 shared SIP genes that suggests that they have elevated expression levels in hematopoietic cell types [17]compared to the gene expression in various other tissues (Methods, Figure S5). We find similar trends when comparing the gene expression of cell type-specific SIP genes to the expression in various other tissues (Figure S6).

### Partitioned Heritability for Cell Type-Specific SIP Genes using GWAS Summary Statistics

We leveraged linkage disequilibrium score regression [30] (LDSC) using the cell type-specific SIP PIRs to partition the SNP heritability using trans-ethnic GWAS summary statistics of 15 blood cell traits [31] (Methods). Enrichment scores and corresponding *p*-values for each cell type and blood cell trait are displayed in Figure S7 and Figure 6. Erythrocyte-specific SIPs are significantly enriched for red blood cell related traits including MCH, MCHC, MCV, RBC and RDW. Further, megakaryocyte-specific SIPs are significantly enriched for PLT, naive CD4 T-cell-specific SIPs are significantly enriched for LYM, and neutrophil-specific SIPs are significantly enriched for NEU and WBC. These results all show expected trait enrichments for each cell type. We also notice some less expected enrichments between erythrocyte-specific SIPs and NEU, as well as between neutrophil-specific SIP genes and MCH, for example. While macrophage/monocyte-specific SIPs are not enriched for white blood cell related traits (including monocyte counts), this may be due to the small number of macrophage/monocyte-specific SIPs (107) relative to the larger number of naive CD4 T-cell-and neutrophil-specific SIPs (283 and 274, respectively). When considering the PIRs of all SIPs, rather than only cell type specific SIP PIRs, macrophage/monocyte SIPs are significantly enriched for MONO and WBC (Figure S8).

**Figure 6.**
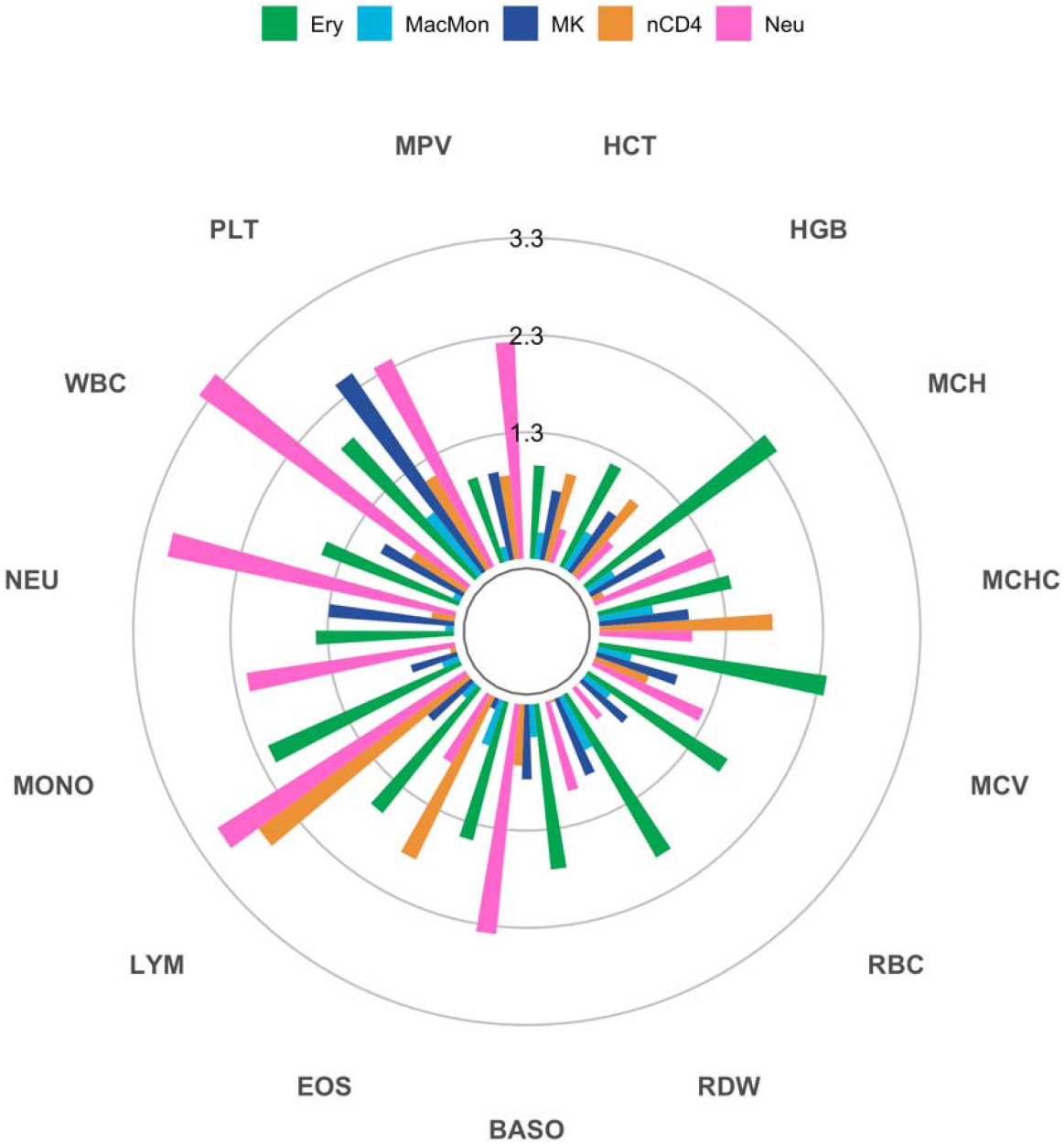
Partitioned heritability for blood cell traits shows enrichment between cell type-specific SIPs and relevant traits. Enrichment score p-values (-log10 scale) for cell type-specific genes and 15 blood cell traits. Bars passing the inner ring (1.3) correspond to statistically significant enrichment scores (*p*-value < 0.05). See Figure S7 for enrichment scores. (Cell types: Ery = erythrocytes; MacMon = macrophages/monocytes; MK = megakaryocytes; nCD4 = naive CD4 T-cells; Neu = neutrophils. Blood cell traits: HCT = Hematocrit; HGB = Hemoglobin; MCH = Mean Corpuscular Hemoglobin; MCHC = MCH Concentration; MCV = Mean Corpuscular Volume; RBC = Red Blood Cell Count; RDW = RBC Distribution Width; BASO = Basophil Count; EOS = Eosinophil Count; LYM = Lymphocyte Count; MONO = Monocyte Count; NEU = Neutrophil Count; WBC = White Blood Cell Count; PLT = Platelet Count; MPV = Mean Platelet Volume)

## Discussion

Hi-C has been widely adopted to study chromatin spatial organization. pcHi-C, a derivative of the Hi-C technology, enables the study of the promoter interactome, specifically. Importantly, recent studies have demonstrated the ability of pcHi-C analysis to link non-coding variants to their target genes.

By analyzing pcHi-C data, we catalogue super-interactive promoters (SIPs) in five blood cell types and present characteristics and analysis of SIPs in blood cell lineages. The characteristics of SIPs identified in blood cell lineages are consistent with those described of SIPs identified in the brain cortex [8], including enrichment for key blood lineage-specific genes, cell type specificity for most identified SIPs, and cell type-specific SIP enrichment in cells with higher expression of the regulated genes. We also demonstrate that SIPs share common properties across cell types, but align with cell type-specific genes. In our analyses, we find that SIPs’ regulatory networks are more likely to overlap with relevant GWAS variants and ATAC-seq peaks than non-SIP regulatory networks. We further find that cell type-specific SIP genes show enriched heritability in blood cell trait GWAS summary statistics. In conjunction with other functional genomic data, we hypothesize that SIPs in relevant hematopoietic cell types can help identify GWAS variant target genes.

Now that many blood cell lineage SIPs have been identified, a logical next step would be to disrupt SIPs or SIP PIRs and evaluate the effects on hematopoiesis. SIPs driven by *few* super strong interactions vs *many* significant (not necessarily all strong) interactions will have different implications for the design and prioritization of functional experiments. In our study, the latter seems to be the norm. Most SIPs are linked to multiple regulatory regions (as opposed to just having a few very strong interactions). These multiple regulatory regions are likely key for orchestrating fine transcriptional control of genes with SIPs. Multiple regulatory regions may also provide a level of “redundancy”, ensuring that even in the presence of an enhancer-disrupting genetic variant, appropriate transcriptional regulation can occur for important genes in a given hematopoietic cell type. Many key GWAS loci show allelic heterogeneity, with multiple rare and common variants (both coding and noncoding) impacting gene regulation (for example, at the *MPL* or *JAK2* locus for platelet traits [31–33]. Particularly for SIPs, genetic or epigenetic perturbations of one of these many putative regulatory regions (some of which may be tagged by statistically distinct GWAS SNPs) may be compensated for by other regulatory regions in the orchestra, leading to no apparent effect *in vitro* even when the perturbed region is functional in its native context. Researchers should consider this limitation when prioritizing loci and interpreting functional validation experiment results and may want to consider approaches that genetically or epigenetically edit multiple variants or regulatory regions simultaneously [34]. Cell type specificity of SIPs and their PIRs should also be considered in linking GWAS variants to genes and in design of functional experiments.

The success of SIP characterization in neuronal, and now hematopoietic lineages, suggests the value of cataloguing SIPs in other cell types and incorporating those SIPs with results of GWAS analysis for relevant traits. It would also be interesting to examine condition-specific SIPs, such as different molecular environments triggered by drugs, toxic chemicals, diet, or stress, in various cell types. Doing so would allow for investigation on how gene expression varies in a cell-type specific manner under different environmental conditions. In addition, future work may involve exploring the relationship between super PIRs and super enhancers. Further experimental work to validate the cell type-specific SIP genes and the connection of these genes to corresponding blood cell traits will be required, but many attractive candidates have been identified through our systematic evaluation of promoters and their interacting regulatory regions in hematopoietic cell types.

## Conclusions

We identified 808-1287 SIPs from major blood cell types, corresponding to 1,093-1,752 SIP genes, among which 125-386 are cell type specific. These SIPs and SIP genes in blood cells will be valuable not only for studying hematological traits but for many complex phenotypes. SIPs manifest significant differences from non-SIPs in at least four aspects: (1) promoter-interacting regions (PIRs) of SIPs are more likely to overlap with cell type-specific open chromatin regions; (2) SIP PIRs, compared to PIRs for non-SIPs, are enriched with GWAS variants associated with relevant hematological traits; (3) SIP genes tend to express at a higher level in the corresponding cell type; (4) SIP genes show enriched heritability of relevant blood cell traits.

We provide mechanistic hypotheses regarding the formation of SIPs. To be identified as a SIP, a promoter can be driven by few super strong interactions or many significant (not necessarily all strong) interactions. Importantly, we find that the latter seems to be the norm. This finding sheds light regarding the formation of SIPs: to ensure the expression level of some critical gene (here a SIP gene), multiple regulatory regions are likely key for orchestrating fine transcriptional control. These multiple regulatory regions provide a level of “redundancy”, ensuring that even in the presence of genetic variant(s) that disrupt some enhancer(s), appropriate transcriptional regulation can still be maintained in a given hematopoietic cell type. This finding also has important implications for the interpretation and functional follow-up of hundreds of thousands of GWAS findings. These multiple regulatory regions for one SIP gene help explain multiple independent GWAS signals at one locus. We provide concrete examples, including the *EPHB3* locus associated with platelet count where we present three distinct regulatory regions overlapping a total of 8 independent SNPs. In addition, due to the potential redundancy, functional experiments may also need to consider disrupting multiple regulatory regions simultaneously rather than individually to observe palpable effects. In summary, we believe our work presents important findings governing the orchestrated transcriptional control in blood lineage cell types, and provides valuable insights and resources for the interpretation and follow-up of GWAS findings of many complex traits.

## Methods

### Cell Types

There are eight hematopoietic cell types in the pcHi-C data [13]: M0 macrophage, M1 macrophage, M2 macrophage, monocyte, neutrophil, erythrocyte, naive CD4 T-cell, and megakaryocyte. Since monocytes circulate in the blood and exist in tissues as macrophages in their mature form, we grouped the monocytes with the three macrophage types (by taking the average of the gene expression in BLUEPRINT [17] and the CHiCAGO [26] scores in pcHi-C data) to form one group. Thus, we focus on five cell types throughout this paper.

### Definition of Cell Type-Specific versus Shared Genes

We classified genes as cell type-specific or shared via the Shannon entropy across the five cell types. Gene expression data was downloaded from BLUEPRINT [17]. Since this gene expression is calculated by MMSEQ, we took exponentials so that transcript quantification was comparable to RPKM. For each gene, we calculated the normalized gene expression as the gene expression in one cell type divided by the sum of the gene’s expression across all five cell types. Next, we calculated the entropy (defined as the distance to log2(*K*), where *K*=5 is the number of cell types) using the relative gene expression across cell types. We defined cell type-specific genes as those with entropy > 0.5 and gene expression > 1, in the respective cell type, and shared genes across cell types as those with entropy < 0.1. Approximately 534-1,814 genes are cell type-specific, depending on cell type (Figure S1f), and 1,476 genes meet the shared gene criteria.

### Defining SIPs

We first calculated the cumulative interaction scores for each promoter-containing anchor bin (bait) in the pcHi-C data [13], in each cell type. For each bait, the cumulative interaction score is the sum of the CHiCAGO scores of significant interactions (CHiCAGO score >= 5, as informed by Cairns et al. [26]). Interactions with CHiCAGO score < 5 are not included in the cumulative interaction score. For each cell type, we ranked the cumulative interaction scores. By finding the inflection point of the ranked baits, we defined super interactive promoters (SIPs) as those baits with extremely high cumulative interaction scores. This approach is similar to how super-enhancers are defined [35]. SIPs are approximately the top 7.5% of cumulative interaction scores.

### SIP PIRs Overlap with Relevant GWAS Variants

In each cell type, for every SIP, we determined if at least one PIR overlapped with a relevant blood cell trait variant, using summary statistics from the latest two GWAS studies on blood cell traits, including GWAS variants identified in European samples [33] as well as non-European and trans-ethnic analyses [31]. Phenotypes (i.e., relevant traits) considered for each cell type are as follows: any red blood cell trait (HCT, HGB, MCH, MCHC, RBC, RDW) for erythrocytes, MONO or WBC for macrophages/monocytes, PLT or MPV for megakaryocytes, LYM or WBC for naive CD4 T-cells, and NEU or WBC for neutrophils. Next, for each cell type, we randomly sampled non-SIPs (where *n* sampled is the number of SIPs in the respective cell type) and determined if at least one PIR overlapped with a relevant variant. This sampling procedure was repeated 100 times, and the median number of non-SIPs with a PIR overlapping a relevant variant was recorded. Fisher’s exact test was then used to compute odds ratios and 95% confidence intervals for the odds of a SIP with variant overlap compared to a non-SIP with variant overlap. The same procedure was conducted for cell type-specific SIPs.

To construct SIP subnetworks, we only considered the conditionally independent GWAS variants from Vuckovic et al. [33]. Consequently, each SIP subnetwork has PIRs that each overlap with a relevant statistically independent variant, as well as a cell type-specific ATAC-seq peak [27]. We identify SIP subnetworks for each of the following cell type/phenotype combinations: erythrocytes (HCT (2), HGB (2), MCH (7), MCHC (3), RBC (4), RDW (11)), macrophages/monocytes (MONO (5), WBC (1)), megakaryocytes (PLT (14), MPV (10)), and naive CD4 T-cells (LYM (15), WBC (2)). When removing the constraint of PIR overlapping with ATAC-seq data for neutrophil SIPs, as it is unavailable, we identify neutrophil SIP subnetworks for NEU (16) and WBC (22).

### Fold Enrichment Test for Highly Expressed Genes Among Genes with SIPs

Gene expression was ranked from highest (1st) to lowest (5th) quintile in each cell type. For each cell type, we calculated the proportion of SIP genes with rank *r* out of the total number of genes with rank *r*. Fold enrichment was then calculated relative to the group with the lowest gene expression (5th) and the significance level was obtained through a Chi-square test for proportions (for each cell type).

### Comparing Gene expression Levels in Shared and Cell Type-Specific SIP Genes

We downloaded gene expression data for all tissues from the GTEx portal [36]. For comparison to our blood cell types of interest, we used exponentiated BLUEPRINT gene expression [17] for erythrocytes, macrophages/monocytes, megakaryocytes, naive CD4 T-cells, and neutrophils. For each of the shared SIP genes, we computed the mean gene expression across all five blood cell types and the mean gene expression across all other tissues (non-blood cells). Next, we partitioned the shared SIP genes into percentiles based on the ranked mean gene expressions in blood cells (Figure S4a-b), and the ranked mean gene expressions in other tissues (Figure S4c-d). We followed a similar computational process for the cell type-specific SIP genes. For each set of cell type-specific SIP genes, we partitioned the genes into percentiles based on the ranked gene expression in the respective cell type (Figure S5a-b), and the ranked mean gene expressions in other tissues (Figure S5c-d).

### Partitioned Heritability for Cell Type-Specific SIP Genes

We leveraged linkage disequilibrium score regression [30] (LDSC) using the cell type-specific SIP PIRs to partition the SNP heritability for 15 blood cell traits from trans-ethnic GWAS summary statistics [31]. LDSC jointly models 75 baselines annotations consisting of coding, UTR, promoter, and intron regions, histone marks, DNase I hypersensitive sites, ChromHMM/Segway predictions, regions that are conserved in mammals, super-enhancers, FANTOM5 enhancers, and LD-related annotations (recombination rate, nucleotide diversity CpG content, etc.) that are not specific to any cell type.

## Supplemental Note 1

Neutrophils were excluded from the SIP and ATAC-seq peak analysis. The Ulirsch et al. [27] ATAC-seq data used in this analysis does not include neutrophil ATAC-seq. Chen et al. [37] isolated neutrophils from two healthy donors’ blood. Peak calling of this data (performed by MACS2 narrow peak mode with default parameters –q 0.01 –nomodel –shift 0) resulted in ∼2,000 neutrophil ATAC-seq peaks, which is 1-2 orders of magnitude smaller than expected based on ATAC-seq in other hematopoietic cell types and consistent with the findings of Chen et. al who note that neutrophils have fewer chromatin accessibility peaks than do cell types with comparable sequencing depths and alignment rates. Since neutrophils are terminally differentiated cells with a short lifespan and the accessible chromatin peaks are not associated with usual euchromatin marks, it is possible that ATAC-seq peaks are not enriched or relevant for neutrophil traits. We also note that ATAC-seq data from monocytes was used for analyses involving the macrophage/monocyte SIPs.

## Abbreviations of Blood Cell Traits

HCT: Hematocrit
HGB: Hemoglobin
MCH: Mean Corpuscular Hemoglobin
MCHC: MCH Concentration
MCV: Mean Corpuscular Volume
RBC: Red Blood Cell Count
RDW: RBC Distribution Width
BASO: Basophil Count
EOS: Eosinophil Count
LYM: Lymphocyte Count
MONO: Monocyte Count
NEU: Neutrophil Count
WBC: White Blood Cell Count
PLT: Platelet Count
MPV: Mean Platelet Volume

## Description of Supplemental Material and Additional Files

The supplemental material pdf includes 8 figures and 3 tables. There are three additional Excel files: Additional File 1 details information pertaining to SIPs and SIP genes in each cell type, Additional File 2 details SIP PIR overlaps with GWAS variants, and Additional File 3 details information regarding the SIP subnetworks.

## Declarations

### Availability of data and materials

This study did not generate any data. Data generated as a result of the analyses in this study are included as additional files.

### Competing interests

The authors declare that they have no competing interests.

## Funding

This material is based upon work supported by the National Science Foundation Graduate Research Fellowship Program under Grant No. DGE-1650116, awarded to T.M.L. T.M.L., L.M.R., and Y.L. are partially funded by the National Institutes of Health grant R01 HL129132 (awarded to Y.L.). Y.L. is additionally supported by the National Institutes of Health grants R01 GM105785 and P50 HD103573. The laboratory of V.G.S. received support from the New York Stem Cell Foundation and National Institutes of Health grant R01 DK103794. V.G.S. is a New York Stem Cell Foundation-Robertson Investigator. The project described was also supported by the National Center for Advancing Translational Sciences, National Institutes of Health, through Grant KL2TR002490 (L.M.R.). The content is solely the responsibility of the authors and does not necessarily represent the official views of the NIH. L.M.R. was also funded by T32 HL129982.

## Authors’ contributions

Y.L. and M.H. conceived the study. T.M.L., J.W., Y.H., and Y.Y. performed data analysis. All authors contributed to writing and/or editing the paper and approved the final manuscript.

## Acknowledgements

We would like to acknowledge Jacob C. Ulirsch for processing the ATAC-seq data used in this study.

## Notes

### Competing Interest Statement

The authors have declared no competing interest.

