## Supplemental Materials for "Super interactive promoters provide insight into cell type-specific regulatory networks in blood lineage cell types"

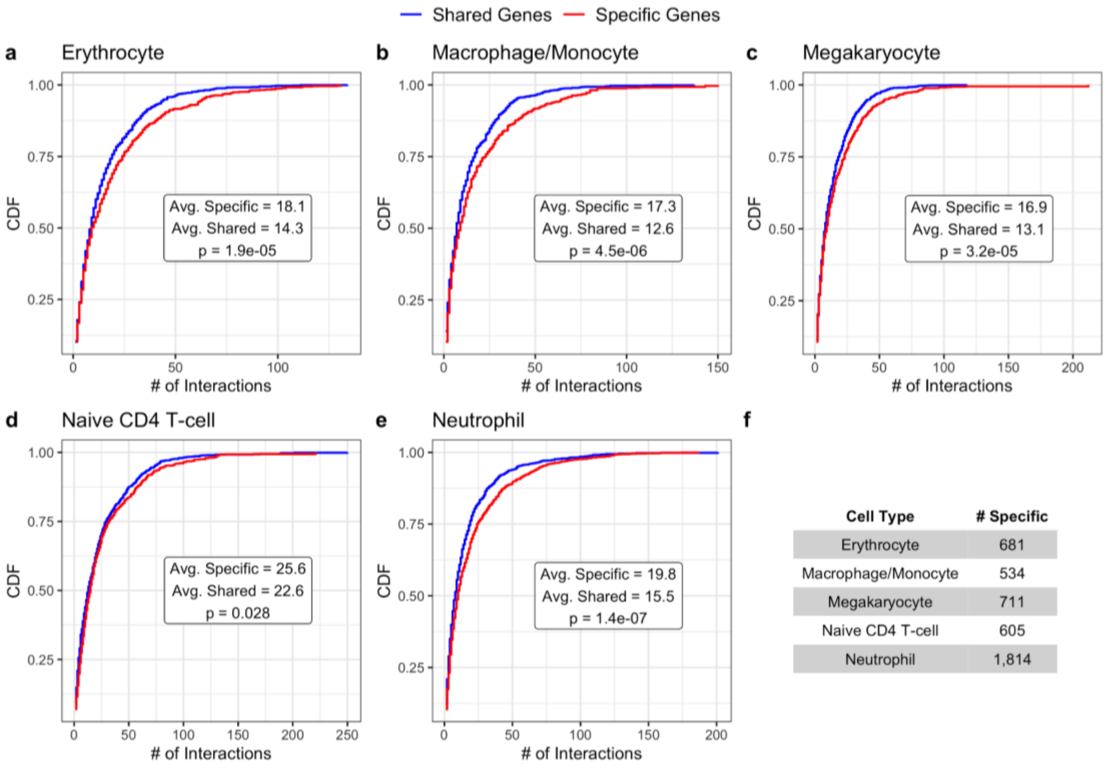

| Cell Type | # Specific |
|---|---|
| Erythrocyte | 681 |
| Macrophage/Monocyte | 534 |
| Megakaryocyte | 711 |
| Naive CD4 T-cell | 605 |
| Neutrophil | 1,814 |

**Figure S1. Cell type-specifically expressed genes exhibit higher levels of chromatin interactivity in the corresponding cell type.** Each panel displays the empirical cumulative distribution function (CDF) of the number of significant pcHi-C interactions for shared versus cell type-specific genes in **(a)** erythrocytes, **(b)** macrophages/monocytes, **(c)** megakaryocytes, **(d)** naive CD4 T-cells, and **(e)** neutrophils. The average number of interactions for specific and shared genes within each cell type is reported along with the corresponding two-sided *t*-test *p*-value. **(f)** The number of specific genes per cell type.

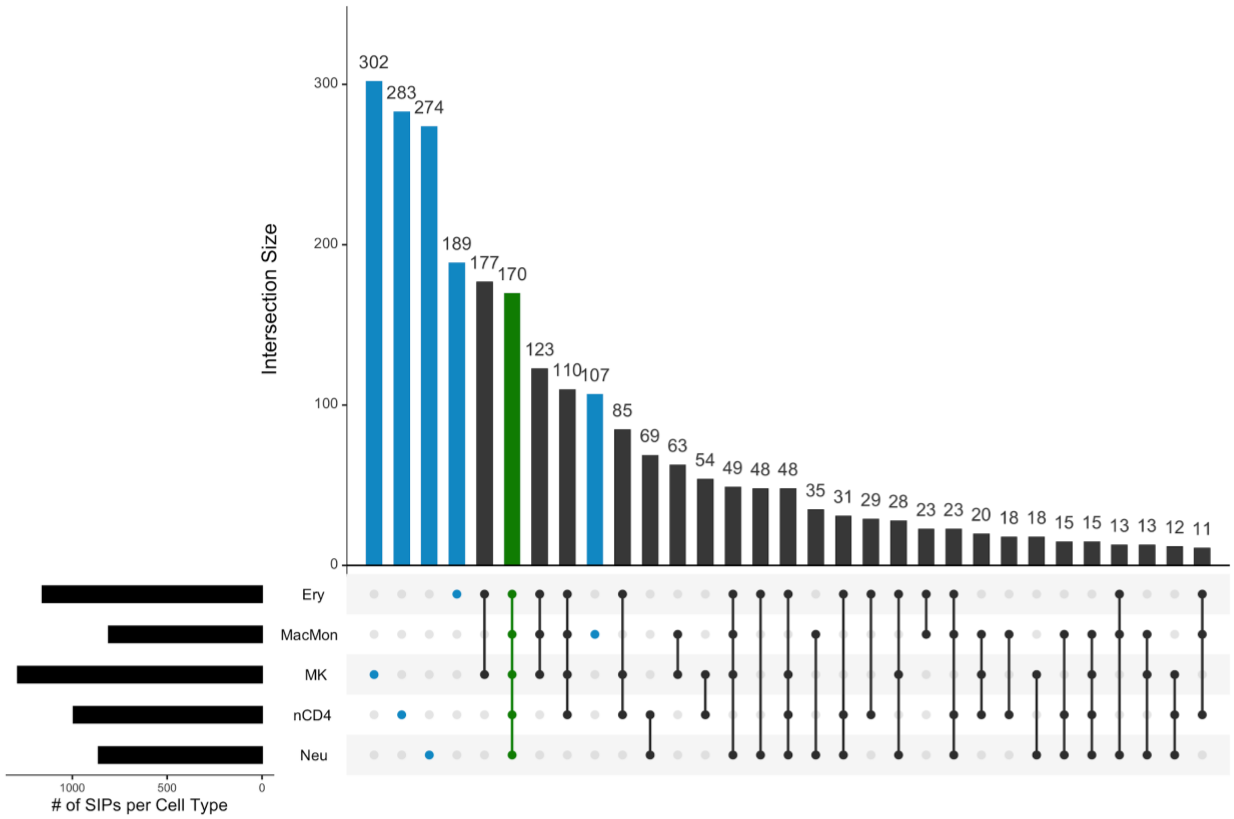

**Figure S2. A majority of SIPs are cell type-specific or shared across all five cell types.** Details of SIPs are shared across cell types (black). Most SIPs, however, are cell-type specific (blue) or common between all five cell type groups (green). (Ery = erythrocytes; MacMon = macrophages/monocytes; MK = megakaryocytes; nCD4 = naive CD4 T-cells; Neu = neutrophils)

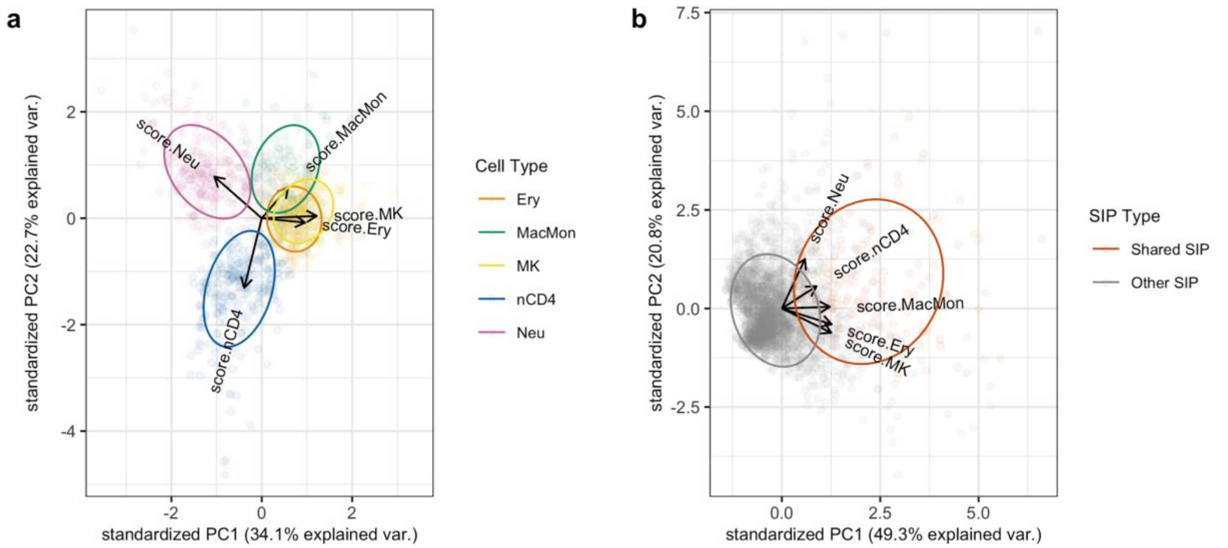

**Figure S3. Principal component analysis (PCA) on cumulative interaction scores reflects expected correlations between SIPs. (a)** PCA on the cumulative interaction scores of cell type-specific SIPs shows most correlation between erythrocyte- and megakaryocyte-specific SIPS, and distinction between those SIPs and the macrophage/monocyte-, naive CD4 T-cell- and neutrophil-specific SIPs (all immune function-related cell types), reflecting known relationships on the hematopoietic tree. **(b)** PCA on the cumulative interaction scores of SIPs, where “Shared” refers to SIPs shared across all five cell types, and “Other” refers to a SIP in at least one cell type.

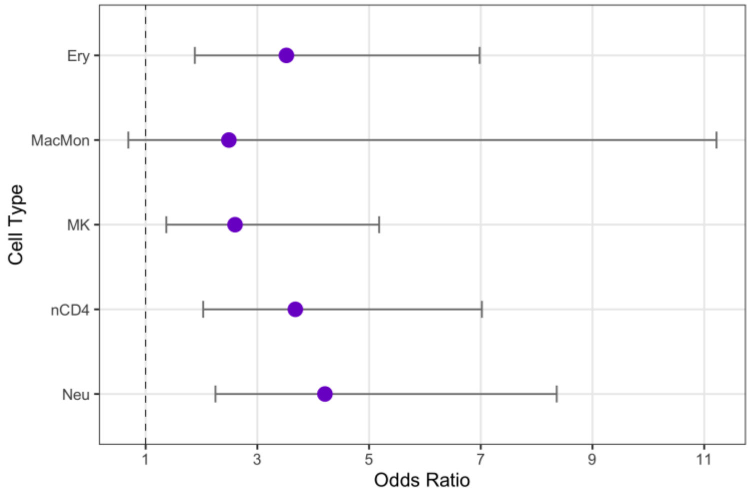

**Figure S4. PIRs of cell type-specific SIPs have greater odds of overlapping GWAS variants.** In each cell type, SIPs have greater odds of having at least one PIR overlap with a relevant GWAS variant, compared to non-SIPs. Odds ratio estimates (purple dots) and corresponding 95% confidence intervals are shown. The large confidence interval seen in macrophages/monocytes is due to the relatively few number of cell type-specific SIPs.

**Figure S5. Shared SIP genes have elevated expression levels in hematopoietic cell types.** Violin plots showing the distribution of gene expression of shared SIP genes in various tissues as well as the five blood cell types. **(a)** Shared SIP genes with the top 10% of expression among blood cells. **(b)** Shared SIP genes with the top 10-20% of expression among blood cells. **(c)** Shared SIP genes with the top 10% of expression among other tissues (non-blood cells). **(d)** Shared SIP genes with the top 10-20% of expression among other tissues (non-blood cells). (Ery = erythrocytes; MacMon = macrophages/monocytes; MK = megakaryocytes; nCD4 = naive CD4 T-cells; Neu = neutrophils)

*Figure S5 is displayed on the following two pages (pages 5 and 6).*

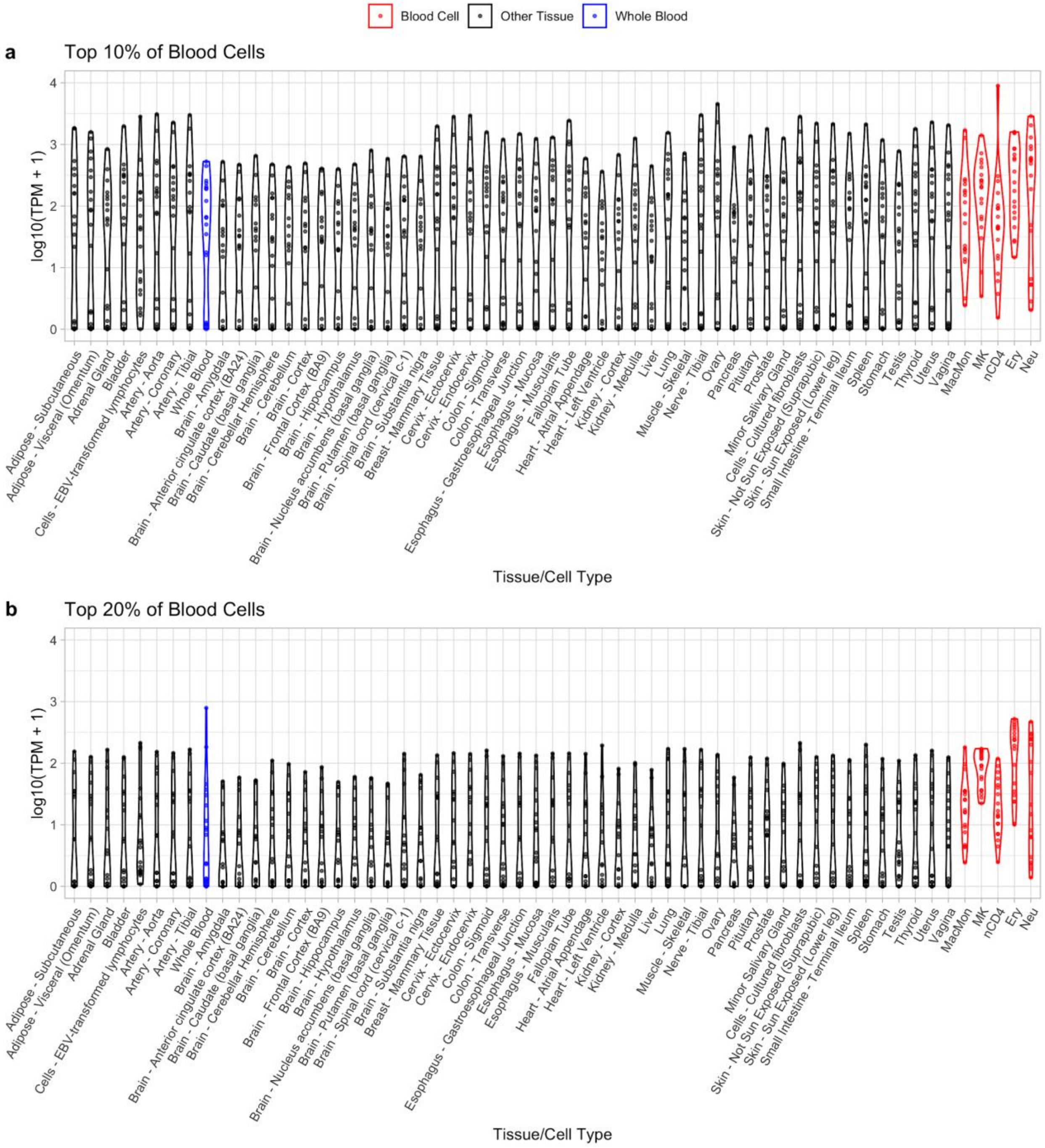
Blood Cell
Other Tissue
Whole Blood
a
Top 10% of Blood Cells
log10(TPM + 1)
0
1
2
3
4
Adipose - Subcutaneous
Adipose - Visceral (Omentum)
Adrenal Gland
Bladder
Cells - EBV-transformed lymphocytes
Artery - Aorta
Artery - Coronary
Artery - Tibial
Whole Blood
Brain - Amygdala
Brain - Anterior cingulate cortex (BA24)
Brain - Caudate (basal ganglia)
Brain - Cerebellar Hemisphere
Brain - Cerebellum
Brain - Cortex
Brain - Frontal Cortex (BA9)
Brain - Hippocampus
Brain - Hypothalamus
Brain - Nucleus accumbens (basal ganglia)
Brain - Putamen (basal ganglia)
Brain - Spinal cord (cervical c-1)
Brain - Substantia nigra
Breast - Mammary Tissue
Cervix - Ectocervix
Cervix - Endocervix
Colon - Sigmoid
Colon - Transverse
Esophagus - Gastroesophageal Junction
Esophagus - Mucosa
Esophagus - Muscularis
Fallopian Tube
Heart - Atrial Appendage
Heart - Left Ventricle
Kidney - Cortex
Kidney - Medulla
Liver
Lung
Muscle - Skeletal
Nerve - Tibial
Ovary
Pancreas
Pituitary
Prostate
Minor Salivary Gland
Cells - Cultured fibroblasts
Skin - Not Sun Exposed (Suprapubic)
Skin - Sun Exposed (Lower leg)
Small Intestine - Terminal Ileum
Spleen
Stomach
Testis
Thyroid
Uterus
Vagina
MacMon
MK
nCD4
Ery
Neu
Tissue/Cell Type
b
Top 20% of Blood Cells
log10(TPM + 1)
0
1
2
3
4
Adipose - Subcutaneous
Adipose - Visceral (Omentum)
Adrenal Gland
Bladder
Cells - EBV-transformed lymphocytes
Artery - Aorta
Artery - Coronary
Artery - Tibial
Whole Blood
Brain - Amygdala
Brain - Anterior cingulate cortex (BA24)
Brain - Caudate (basal ganglia)
Brain - Cerebellar Hemisphere
Brain - Cerebellum
Brain - Cortex
Brain - Frontal Cortex (BA9)
Brain - Hippocampus
Brain - Hypothalamus
Brain - Nucleus accumbens (basal ganglia)
Brain - Putamen (basal ganglia)
Brain - Spinal cord (cervical c-1)
Brain - Substantia nigra
Breast - Mammary Tissue
Cervix - Ectocervix
Cervix - Endocervix
Colon - Sigmoid
Colon - Transverse
Esophagus - Gastroesophageal Junction
Esophagus - Mucosa
Esophagus - Muscularis
Fallopian Tube
Heart - Atrial Appendage
Heart - Left Ventricle
Kidney - Cortex
Kidney - Medulla
Liver
Lung
Muscle - Skeletal
Nerve - Tibial
Ovary
Pancreas
Pituitary
Prostate
Minor Salivary Gland
Cells - Cultured fibroblasts
Skin - Not Sun Exposed (Suprapubic)
Skin - Sun Exposed (Lower leg)
Small Intestine - Terminal Ileum
Spleen
Stomach
Testis
Thyroid
Uterus
Vagina
MacMon
MK
nCD4
Ery
Neu
Tissue/Cell Type

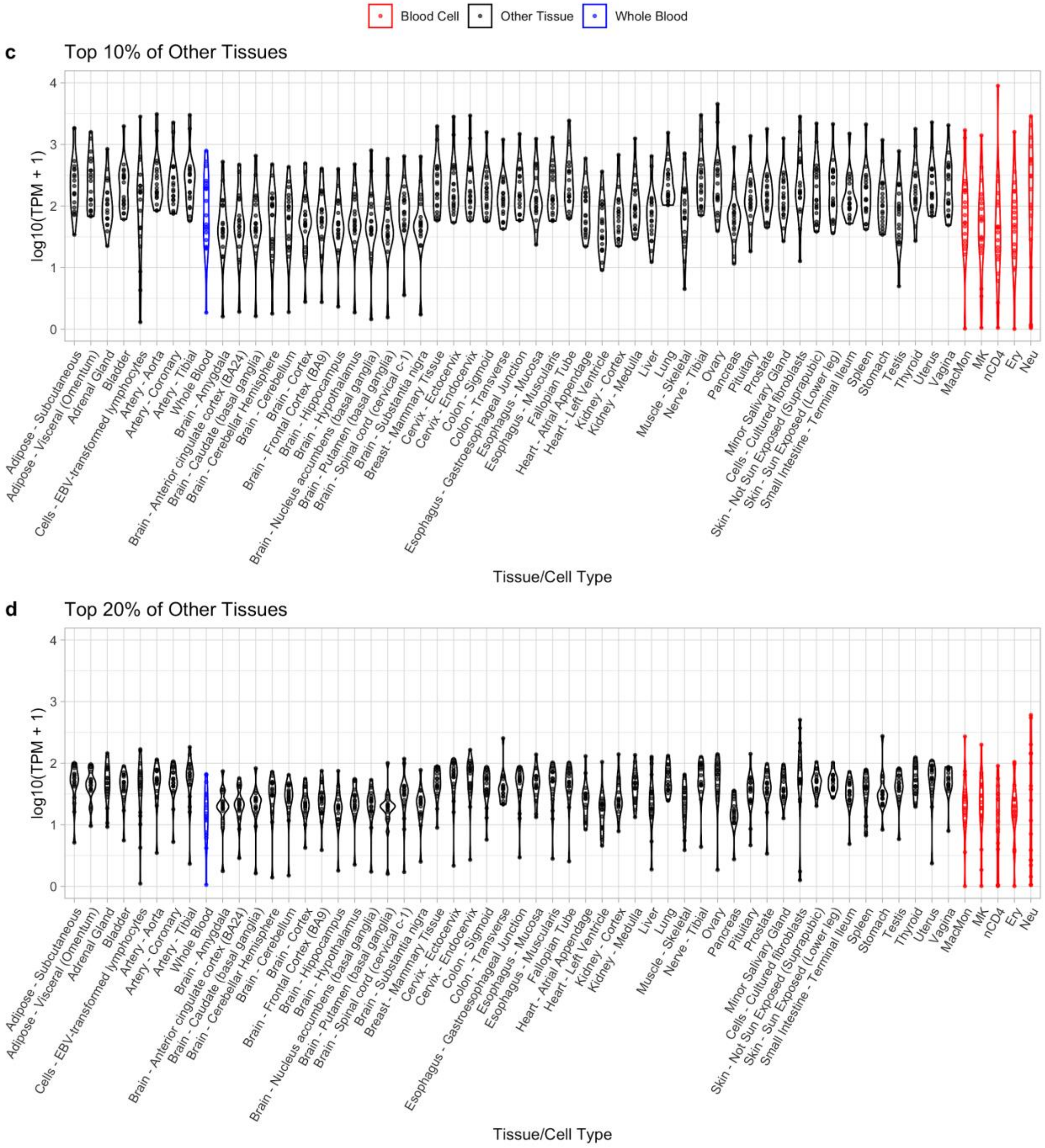
Blood Cell
Other Tissue
Whole Blood
c
Top 10% of Other Tissues
d
Top 20% of Other Tissues
log10(TPM + 1)
0
1
2
3
4
Tissue/Cell Type
Adipose - Subcutaneous
Adipose - Visceral (Omentum)
Adrenal Gland
Bladder
Cells - EBV-transformed lymphocytes
Artery - Aorta
Artery - Coronary
Artery - Tibial
Whole Blood
Brain - Amygdala
Brain - Anterior cingulate cortex (BA24)
Brain - Caudate (basal ganglia)
Brain - Cerebellar Hemisphere
Brain - Cerebellum
Brain - Cortex
Brain - Frontal Cortex (BA9)
Brain - Hippocampus
Brain - Hypothalamus
Brain - Nucleus accumbens (basal ganglia)
Brain - Putamen (basal ganglia)
Brain - Spinal cord (cervical c-1)
Brain - Substantia nigra
Breast - Mammary Tissue
Cervix - Ectocervix
Cervix - Endocervix
Colon - Sigmoid
Colon - Transverse
Esophagus - Gastroesophageal Junction
Esophagus - Mucosa
Esophagus - Muscularis
Fallopian Tube
Heart - Atrial Appendage
Heart - Left Ventricle
Kidney - Cortex
Kidney - Medulla
Liver
Lung
Muscle - Skeletal
Nerve - Tibial
Ovary
Pancreas
Pituitary
Prostate
Minor Salivary Gland
Cells - Cultured fibroblasts
Skin - Not Sun Exposed (Suprapubic)
Skin - Sun Exposed (Lower leg)
Small Intestine - Terminal Ileum
Spleen
Stomach
Testis
Thyroid
Uterus
Vagina
MacMon
MK
nCD4
Ery
Neu

**Figure S6. Neutrophil specific SIP genes have elevated expression levels in neutrophils.** Violin plots showing the distribution of gene expression of neutrophil-specific SIP genes in various tissues as well as the five blood cell types. The cell type-specific SIP genes for erythrocytes, macrophages/monocytes, and naive CD4 T-cells show similar trends. **(a)** Neutrophil-specific SIP genes with the top 10% of neutrophils expression. **(b)** Neutrophil-specific SIP genes with the top 10-20% of neutrophils expression. **(c)** Neutrophil-specific SIP genes with the top 10% of expression among other tissues (non-blood cells). **(d)** Neutrophil-specific SIP genes with the top 10-20% of expression among other tissues (non-blood cells). (Ery = erythrocytes; MacMon = macrophages/monocytes; MK = megakaryocytes; nCD4 = naive CD4 T-cells; Neu = neutrophils)

*Figure S6 is displayed on the following two pages (pages 8 and 9).*

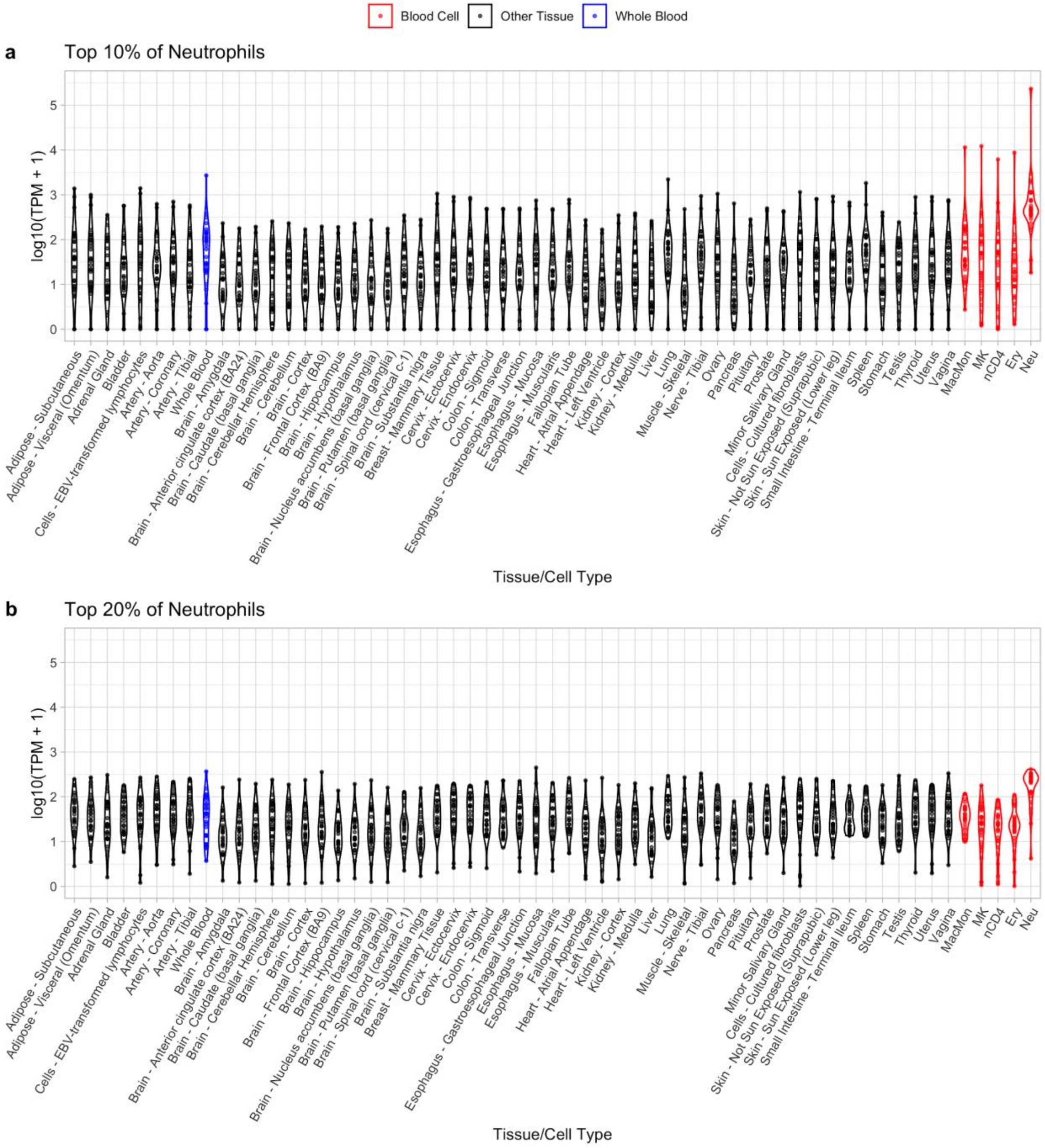
Blood Cell
Other Tissue
Whole Blood
a
Top 10% of Neutrophils
b
Top 20% of Neutrophils
log10(TPM + 1)
Tissue/Cell Type
Adipose - Subcutaneous
Adipose - Visceral (Omentum)
Adrenal Gland
Bladder
Cells - EBV-transformed lymphocytes
Artery - Aorta
Artery - Coronary
Artery - Tibial
Whole Blood
Brain - Amygdala
Brain - Anterior cingulate cortex (BA24)
Brain - Caudate (basal ganglia)
Brain - Cerebellar Hemisphere
Brain - Cerebellum
Brain - Cortex
Brain - Frontal Cortex (BA9)
Brain - Hippocampus
Brain - Hypothalamus
Brain - Nucleus accumbens (basal ganglia)
Brain - Putamen (basal ganglia)
Brain - Spinal cord (cervical c-1)
Brain - Substantia nigra
Breast - Mammary Tissue
Cervix - Ectocervix
Cervix - Endocervix
Colon - Sigmoid
Colon - Transverse
Esophagus - Gastroesophageal Junction
Esophagus - Mucosa
Esophagus - Muscularis
Fallopian Tube
Heart - Atrial Appendage
Heart - Left Ventricle
Kidney - Cortex
Kidney - Medulla
Liver
Lung
Muscle - Skeletal
Nerve - Tibial
Ovary
Pancreas
Pituitary
Prostate
Minor Salivary Gland
Cells - Cultured fibroblasts
Skin - Not Sun Exposed (Suprapubic)
Skin - Sun Exposed (Lower leg)
Small Intestine - Terminal Ileum
Spleen
Stomach
Testis
Thyroid
Uterus
Vagina
MacMon
MK
nCD4
Ery
Neu

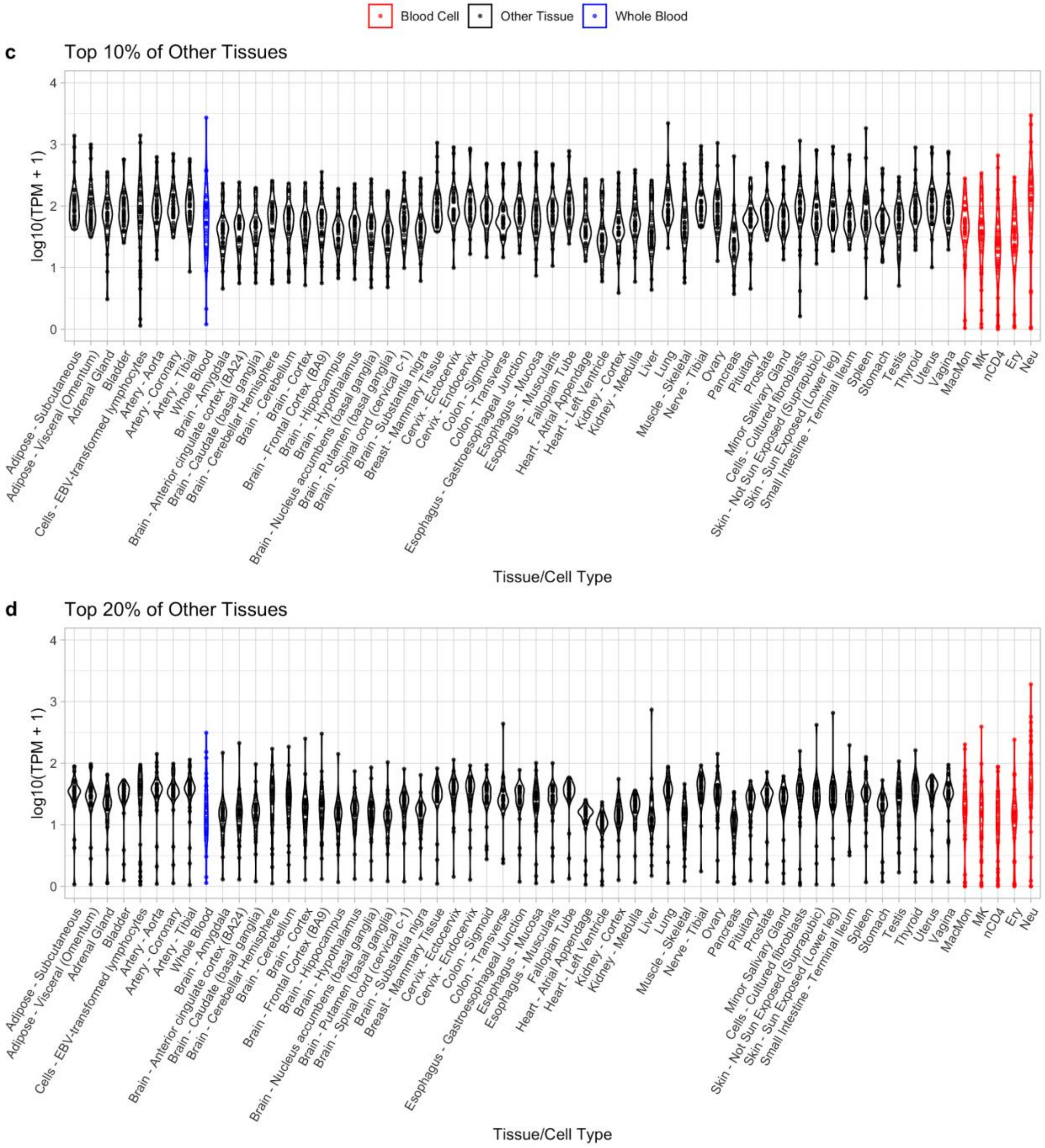

Blood Cell
Other Tissue
Whole Blood
c
Top 10% of Other Tissues
log10(TPM + 1)
0
1
2
3
4
Adipose - Subcutaneous
Adipose - Visceral (Omentum)
Adrenal Gland
Bladder
Cells - EBV-transformed lymphocytes
Artery - Aorta
Artery - Coronary
Artery - Tibial
Whole Blood
Brain - Amygdala
Brain - Anterior cingulate cortex (BA24)
Brain - Caudate (basal ganglia)
Brain - Cerebellar Hemisphere
Brain - Cerebellum
Brain - Cortex
Brain - Frontal Cortex (BA9)
Brain - Hippocampus
Brain - Hypothalamus
Brain - Nucleus accumbens (basal ganglia)
Brain - Putamen (basal ganglia)
Brain - Spinal cord (cervical c-1)
Brain - Substantia nigra
Breast - Mammary Tissue
Cervix - Ectocervix
Cervix - Endocervix
Colon - Sigmoid
Colon - Transverse
Esophagus - Gastroesophageal Junction
Esophagus - Mucosa
Esophagus - Muscularis
Fallopian Tube
Heart - Atrial Appendage
Heart - Left Ventricle
Kidney - Cortex
Kidney - Medulla
Liver
Lung
Muscle - Skeletal
Nerve - Tibial
Ovary
Pancreas
Pituitary
Prostate
Minor Salivary Gland
Cells - Cultured fibroblasts
Skin - Not Sun Exposed (Suprapubic)
Skin - Sun Exposed (Lower leg)
Small Intestine - Terminal Ileum
Spleen
Stomach
Testis
Thyroid
Uterus
Vagina
MacMon
MK
nCD4
Ery
Neu
Tissue/Cell Type
d
Top 20% of Other Tissues
log10(TPM + 1)
Tissue/Cell Type

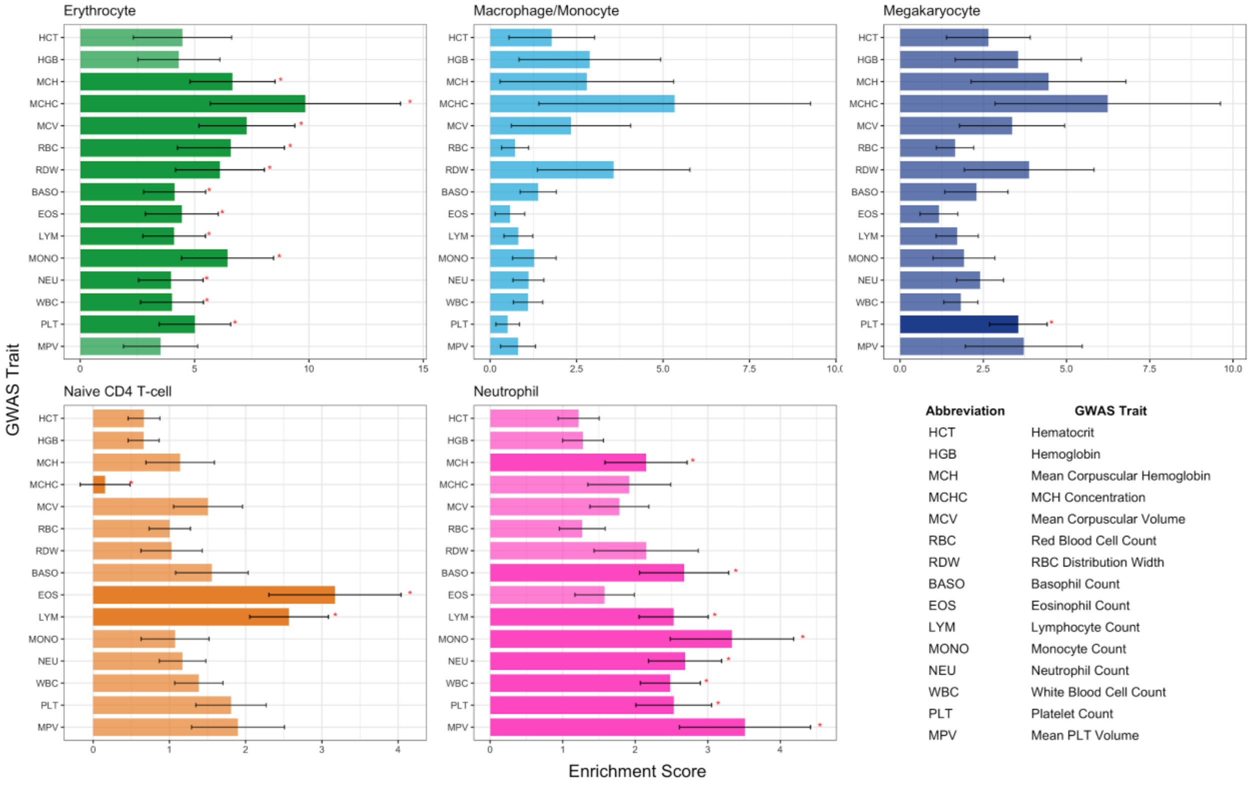

**Figure S7. Partitioned SNP heritability for blood cell traits - cell type-specific SIPs.** Enrichment scores for cell type-specific SIPs and 15 blood cell traits. (* denotes statistically significant enrichment score ($p < 0.05$); Ery = erythrocytes; MacMon = macrophages/monocytes; MK = megakaryocytes; nCD4 = naive CD4 T-cells; Neu = neutrophils)

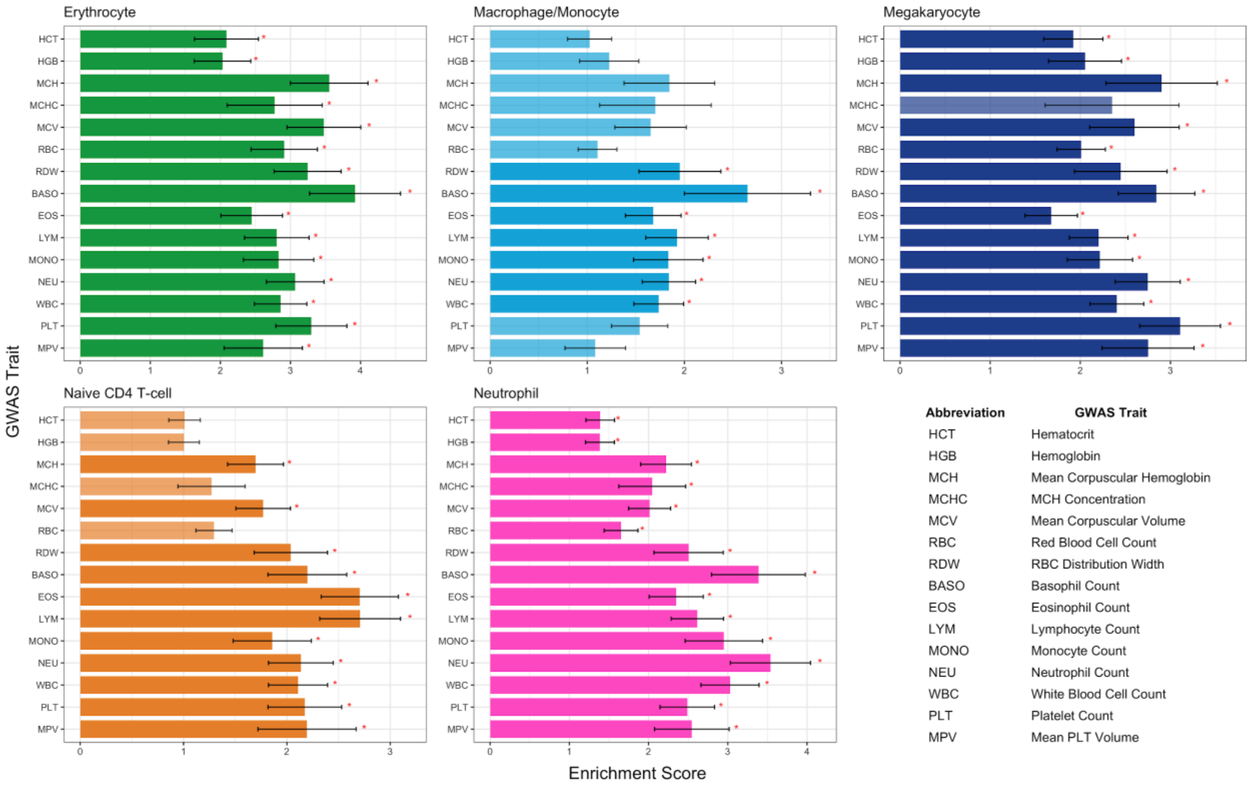

**Figure S8. Partitioned SNP heritability for blood cell traits - all SIPs.** Enrichment scores for SIPs and 15 blood cell traits. (* denotes statistically significant enrichment score ($p$ < 0.05); Ery = erythrocytes; MacMon = macrophages/monocytes; MK = megakaryocytes; nCD4 = naive CD4 T-cells; Neu = neutrophils)

**Table S1. Total counts of cell type-specific SIPs and SIP genes as well as total counts of all SIPs and SIP genes, in each cell type.** The total number of SIPs and SIP genes shared across all five cell types is also reported. Percent refers to the percent of total SIPs or SIP genes that are cell type-specific.

| **SIPs** | | | |
|---|---|---|---|
| | # Specific | # Total | Percent |
| Erythrocyte | 189 | 1,157 | 16.3% |
| Macrophage/Monocyte | 107 | 808 | 13.2% |
| Megakaryocyte | 302 | 1,287 | 23.4% |
| Naive CD4 T-cell | 283 | 993 | 28.5% |
| Neutrophil | 274 | 861 | 31.8% |
| Shared | *NA* | 170 | *NA* |
| **SIP Genes** | | | |
| | # Specific | # Total | Percent |
| Erythrocyte | 251 | 1,615 | 15.6% |
| Macrophage/Monocyte | 125 | 1,093 | 11.4% |
| Megakaryocyte | 385 | 1,752 | 22.0% |
| Naive CD4 T-cell | 386 | 1,393 | 27.7% |
| Neutrophil | 284 | 1,201 | 32.0% |
| Shared | *NA* | 234 | *NA* |

**Table S2. SIPs interact with more super promoter-interacting regulatory regions (super PIRs) than non-SIPs.** Details corresponding to Figure 4. For each cell type, the number of promoter baits interacting with a super PIR or typical PIR are reported, along with the corresponding ratio and Chi-square p-value. Median PIR scores for each cell type and promoter bait type are also reported, along with the corresponding Wilcoxon p-value for the difference in distribution between SIPs and non-SIPs. (Ery = erythrocytes; MacMon = macrophages/monocytes; MK = megakaryocytes; nCD4 = naive CD4 T-cells; Neu = neutrophils)

| | Bait Type | # Super PIR | # Typical PIR | Ratio | Chi-sq p-value | Median PIR Score | Wilcoxon p-value |
|---|---|---|---|---|---|---|---|
| **Ery** | | | | | | | |
| | SIP | 969 | 188 | 0.84 | 2.7e-91 | 6 | 1.6e-119 |
| | non-SIP | 6,924 | 6,188 | 0.53 | | 4 | |
| **MacMon** | | | | | | | |
| | SIP | 603 | 205 | 0.75 | 3.2e-35 | 5 | 1.7e-50 |
| | non-SIP | 6,550 | 6,011 | 0.52 | | 4 | |
| **MK** | | | | | | | |
| | SIP | 955 | 332 | 0.74 | 2.7e-52 | 5 | 2.9e-68 |
| | non-SIP | 6,796 | 6,273 | 0.52 | | 4 | |
| **nCD4** | | | | | | | |
| | SIP | 889 | 104 | 0.90 | 7.7e-50 | 7 | 3.2e-69 |
| | non-SIP | 8,311 | 4,146 | 0.67 | | 5 | |
| **Neu** | | | | | | | |
| | SIP | 716 | 145 | 0.83 | 8.6e-83 | 6 | 2.9e-104 |
| | non-SIP | 5,399 | 5,618 | 0.49 | | 3 | |

**Table S3. SIP PIRs overlap with ATAC-seq peak regions.** Details corresponding to Figure 6. For each cell type, the number of SIP PIRs and non-SIP PIRs that overlap with a cell type-specific ATAC-seq peak region are reported, along with the corresponding ratio and Chi-square p-value. The median (Med.) and average (Avg.) number of PIR interactions overlapping an ATAC-seq peak and corresponding t-test *p*-value for the difference between SIP and non-SIP PIRs is also reported. (Ovlp. = Overlap; Ery = erythrocytes; MacMon = macrophages/monocytes; MK = megakaryocytes; nCD4 = naive CD4 T-cells; Neu = neutrophils)

| | Bait Type | # Overlap | # No Overlap | Ratio | Chi-sq p-value | Med. # Ovlp. | Avg. # Ovlp. | t p-value |
|---|---|---|---|---|---|---|---|---|
| **Ery** | | | | | | | | |
| | SIP | 1,118 | 39 | 0.97 | 5.3e-118 | 8 | 9.4 | 1.2e-162 |
| | non-SIP | 8,255 | 4,857 | 0.63 | | 1 | 1.9 | |
| **MacMon** | | | | | | | | |
| | SIP | 798 | 10 | 0.99 | 8.9e-52 | 12 | 12.8 | 2.7e-172 |
| | non-SIP | 9,491 | 3,070 | 0.76 | | 2 | 2.6 | |
| **MK** | | | | | | | | |
| | SIP | 1,285 | 2 | 1.00 | 4.7e-63 | 12 | 13.5 | 6.3e-292 |
| | non-SIP | 10,645 | 2,424 | 0.81 | | 2 | 2.9 | |
| **nCD4** | | | | | | | | |
| | SIP | 992 | 1 | 1.00 | 2.9e-45 | 22 | 23.3 | 2.1e-290 |
| | non-SIP | 10,315 | 2,142 | 0.83 | | 3 | 4.4 | |

### Description of Supplemental Excel Files:

**Additional File 1. Details of SIPs and SIP genes.** Each sheet in the Excel file details the SIPs and SIP genes for each of the five cell types (Ery = erythrocytes; MacMon = macrophages/monocytes; MK = megakaryocytes; nCD4 = naive CD4 T-cells; Neu = neutrophils). The columns *baitID*, *baitChr*, *baitStart*, *baitEnd*, and *baitName* pertain to the promoter bait identifier, gene(s), and bait location (from the Javierre et al.[1] pcHi-C data). The column *intscore* refers to the cumulative interaction score calculated to define SIPs, and the column *rank* reflects the order of SIP baits from largest score to smallest. Note that a SIP bait may have multiple genes and genes may correspond to multiple SIP baits. The column *specific* indicates if the SIP gene is only found in this particular cell type, and the column *shared* indicates if the SIP gene is shared across all five cell types. The next five columns, *SIP.Ery, SIP.MacMon, SIP.MK, SIP.nCD4,* and *SIP.Neu*, indicate whether that SIP is also found in the respective cell type. The column *ENSEMBL_ID* indicates the ENSEMBL ID (GRCh37), if found, for the gene names (*baitName)* provided in the pcHi-C data. The subsequent columns *Ery, MacMon, MK, nCD4,* and *Neu*, report the exponentiated BLUEPRINT gene expression (equivalent to RPKM) data in that cell type, if available.

**Additional File 2. Details of SIP PIR overlap with GWAS Variants.** Each sheet in the Excel file provides information for the SIP PIRs that overlap with a relevant GWAS variant, for each of the five cell types (Ery = erythrocytes; MacMon = macrophages/monocytes; MK = megakaryocytes; nCD4 = naive CD4 T-cells; Neu = neutrophils). The columns *baitID*, *baitChr*, *baitStart*, *baitEnd*, and *baitName* pertain to the promoter bait identifier, gene(s), and bait location (from the Javierre et al.[1] pcHi-C data). The columns *oeStart*, *oeEnd*, *oeID*, and *oeName* pertain to the other end (PIR) location, identifier, and gene(s). The column *specific* indicates if the SIP gene is only found in this particular cell type (i.e., a cell type-specific SIP). The remaining columns pertain to the GWAS variants: *Phenotype, VariantID, rsID, Position* (hg19 base pair location of variant)*, pval, ancestry,* and *data* (Vuckovic et al.[2] or Chen et al.[3]).

**Additional File 3. Details of SIP Subnetworks.** Each sheet in the Excel file details the SIP subnetworks for a cell type and a particular phenotype. The naming convention for each sheet is “CellType_Phenotype”. A table of abbreviations and total SIP subnetworks for the cell type/phenotype combinations can be found on the first sheet, “README”, along with column descriptions.

1. Javierre, B. M. *et al.* Lineage-Specific Genome Architecture Links Enhancers and Non-coding Disease Variants to Target Gene Promoters. *Cell* **167,** 1369–1384.e19 (2016).
2. Vuckovic, D. *et al.* The polygenic and monogenic basis of blood traits and diseases. *Cell* **182,** 1214–1231.e11 (2020).
3. Chen, M.-H. *et al.* Trans-ethnic and Ancestry-Specific Blood-Cell Genetics in 746,667 Individuals from 5 Global Populations. *Cell* **182,** 1198–1213.e14 (2020).
